# KDM7-mediated oxygen sensing reprograms chromatin to enhance hypoxia tolerance in the root

**DOI:** 10.1101/2025.11.24.690241

**Authors:** Daai Zhang, Ximena Chirinos, Alessia Del Chiaro, Vinay Shukla, Alexander Ryder, Ane Del Pozo Beltrán, Sergio Iacopino, Pedro Bota, Dušan Živković, Francesco Fioriti, Yuri Telara, Cara J. Ellison, Filippo Costa, Paul R. Elliott, Federico Giorgi, Beatrice Giuntoli, Emily Flashman, Isidro Abreu, Francesco Licausi

## Abstract

Roots frequently encounter low oxygen (hypoxia) from soil compaction or water saturation and must adapt to this stress. We investigated how root tip cells sense hypoxia and adjust the meristem epigenome to activate genes that promote tolerance and growth under oxygen limitation. In Arabidopsis root tips, hypoxia tolerance was linked to increased trimethylation of histone H3 at lysine 4 (H3K4me3). We also found that group 7 demethylases (KDM7s) are directly inhibited by hypoxia, and that genetic inactivation of KDM7s, like hypoxia, induces expression of genes essential for meristem survival under oxygen deprivation. We propose that KDM7s function as root-specific oxygen sensors that prime and support hypoxia tolerance.

## Introduction

Vascular plants use roots to anchor themselves to the substrate and extend them in search of nutrients. Root apical growth is sustained by a pool of stem cells that continuously divide, elongate and differentiate to generate new tissues for water and mineral uptake. During soil exploration, roots often face reduced oxygen availability (hypoxia) due to diffusion limitations, such as under water saturation, or waterlogging. Notably, the evolutionary emergence of roots in plants coincided with the recruitment of group VII Ethylene Response Factors (ERFVIIs) by the oxygen-sensing N-degron pathway (NDP) for protein degradation (*1*, *2*). This, together with the heightened susceptibility of *erfVII* mutants to waterlogging, highlights the essential role of hypoxia acclimation mechanisms in root survival (*3*).

### Priming of hypoxia tolerance in roots is associated with increased H3K4 trimethylation

Exposure of plants to mild hypoxia has been shown to prepare roots and shoots to withstand more severe oxygen deprivation, a process known as priming (*4–7*). To test whether hypoxia priming relies on the established oxygen-sensing N-degron pathway (NDP), we used *Arabidopsis thaliana* mutants defective in this pathway. The NDP functions through sequential N-terminal modifications of regulatory proteins, including ERFVIIs, leading to their ubiquitination and proteasomal degradation (**Fig. 1A**) (*8*). We compared the survival and growth of wild-type Columbia-0 (Col-0) with NDP mutants *proteolysis6* (*prt6*) and *arginyl transferase1/2* (*ate1/2*) after strict hypoxia (24 h at 0.1% O₂), following either mild hypoxia priming (24 h at 7% O₂) or control conditions (24 h at 21% O₂) (**Fig. 1B**). Unexpectedly, although *prt6* and *ate1/2* mutants showed moderately enhanced survival without priming, pre-exposure to mild hypoxia improved root survival and growth to a similar extent in all genotypes (**Fig. 1B-C**).Since all NDP substrates, including the ERFVIIs, are expected to be stabilised in *prt6* and *ate1/2*, we concluded that a distinct mechanism must exist to prime root hypoxic tolerance.

**Figure 1.**
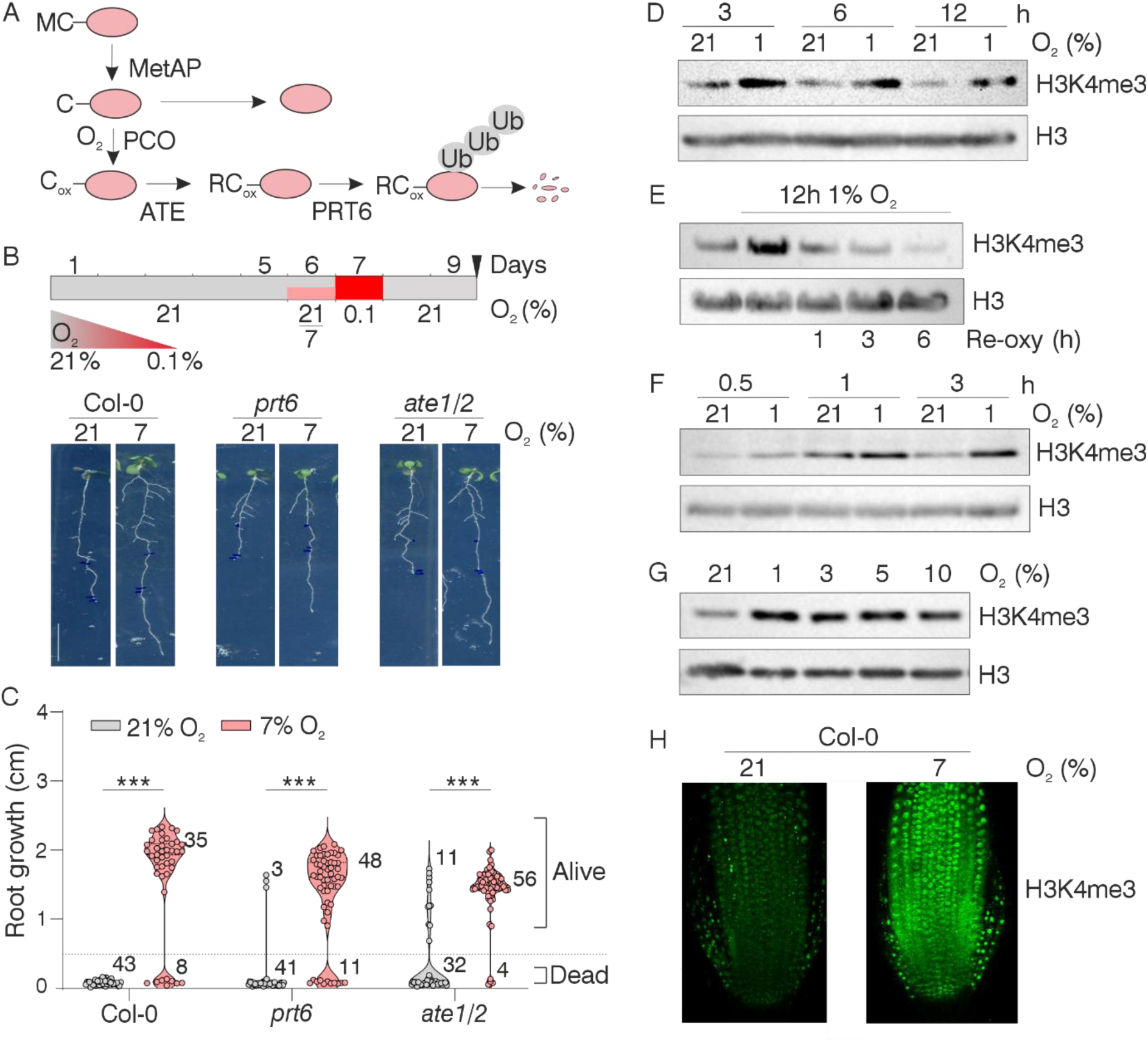
Oxygen-sensitive H3K4 methylation, but not the Cys-N-degron pathway (Cys-NDP), is associated with hypoxia acclimation in root tips. (A) Schematic representation of the Cys-NDP. Proteins containing a cysteine in position 2 can be degraded in an oxygen-dependent manner via the concerted action of methionine aminopeptidase (MetAP), plant cysteine oxidase (PCO), arginyl-tRNA transferase (ATE), and the E3 ubiquitin ligase PRT6. Under hypoxia, these proteins are stabilized and perform their molecular function. (B) A 24 h mild hypoxia pretreatment prior to 24 h of strict hypoxia (0.1% O₂) promotes root growth resumption in wild-type *Arabidopsis thaliana* Col-0 and N-degron pathway mutants *ate1/2* and *prt6* (representative phenotype shown). (C) Quantification of post-hypoxia root growth. Roots exhibiting 0 ± 0.2 mm elongation were classified as “dead.” Mild hypoxia-pretreated (pink) and non-pretreated (grey) “alive” roots were compared using a t-test (*** p ≤ 0.001). (D) Detection of H3K4me3 through immunostaining after 3, 6, and 12 h of hypoxia (1% O₂) or normoxia (21% O₂) in total nuclear proteins extracted from 20-day-old *Arabidopsis* plants. (E) Effect of 1, 3, and 6 h of reoxygenation on H3K4me3 levels following 12 h hypoxia (1% O₂). (F) Detection of H3K4me3 after 0.5, 1, and 3 h of hypoxia (1% O₂) or normoxia (21% O₂) in nuclear extracts from 20-day-old *Arabidopsis* plants. (G) H3K4me3 detection after exposure of 20-day-old *Arabidopsis* plants to varying O₂ concentrations for 12 h. Total H3 is shown in D–G as a loading control. (H) H3K4me3 immunostaining of *Arabidopsis* root tips exposed to mild hypoxia (7% O₂) or normoxia (21% O₂) for 12 h.

In plants, priming against abiotic stresses often occurs through epigenetic reprogramming of chromatin, involving DNA and histone modifications (*9–11*). In mammalian cells, direct oxygen sensing via increased histone methylation establishes a chromatin landscape that facilitates transcriptional regulation by hypoxia-stabilized transcription factors (*12*). We tested whether a similar mechanism operates in plants by probing nuclear protein extracts from Arabidopsis seedlings exposed to aerobic (21% O₂) or hypoxic (1% O₂) conditions for 3, 6, and 12 h. Using antibodies against methylated histone H3 at lysines 4, 9, 27, and 36, we observed fluctuations in H3K9, H3K27, and H3K36 methylation that likely reflect circadian or light-dependent regulation, as previously reported (*13*, *14*) (**Fig. 1D, Supplementary Fig. 1A**). In contrast, H3K4 trimethylation consistently increased under hypoxia, while di- and mono-methylation remained unchanged (**Supplementary Fig. 1B**). We therefore focused further investigation on the role of H3K4 trimethylation in hypoxia responses.

First, we confirmed that H3K4me3 accumulation is reversible upon reoxygenation after 12 h of hypoxia (**Fig. 1E**) and detectable as early as 30 min after hypoxia onset (**Fig. 1F**). We next tested its sensitivity by exposing seedlings to sub-aerobic conditions for 12 h and observed increased H3K4me3 levels at oxygen concentrations between 5–10% (**Fig. 1G**). Given that the hypoxia priming phenotype was observed in roots, we used immunodetection to confirm elevated H3K4me3 in the root tip at 7% O_2_ (**Fig. 1H**). Together, these observations indicate that enhanced root tolerance to severe hypoxia is accompanied by increased H3K4me3, prompting us to investigate whether this chromatin modification directly mediates hypoxia acclimation and how it is mechanistically linked to oxygen availability.

### Genetic inactivation of KDM7s mimics mild hypoxia to promote tolerance in the root tip

Lysine methylation at histones N-terminal tails is mediated by methyltransferases and reversed by demethylases (*15*, *16*). One well-characterized methyltransferase complex is Polycomb Repressive Complex 2 (PRC2), which includes the hypoxia-sensitive VRN2 subunit (*17*). PRC2 specifically targets histone H3 at lysine 27 (H3K27) (*18*). Consistent with this, we observed that increased H3K4me3 levels under hypoxia were not reduced but rather enhanced in the *vrn2* knockout mutant (**Supplementary Fig. 2**). On the other hand, Jumonji C (JmjC) domain-containing histone lysine demethylases (KDMs) are Fe²⁺- and 2-oxoglutarate (2-OG)-dependent dioxygenases that require O₂ as a substrate. These enzymes catalyse hydroxylation of methylated lysines, generating an unstable hydroxymethyl-lysine intermediate that spontaneously decomposes to release formaldehyde, resulting in loss of the methyl group (**Fig. 2A**). The Arabidopsis genome encodes 22 Jmj-C KDMs, grouped in 7 clades based on additional domain architecture (*19*). Different clades preferentially target distinct H3 lysine residues and regulate diverse processes, including flowering time, seed germination, and dehydration responses (*20*). A survey of the literature revealed that H3K4me3 is largely mediated by group 7 KDMs (KDM7A-E), which have also been implicated in root growth control (*21*) (**Supplementary Fig. 3**). Therefore, we decided to test whether genetic inactivation of group 7 KDMs mimics the effects of hypoxia-induced inhibition.

**Figure 2.**
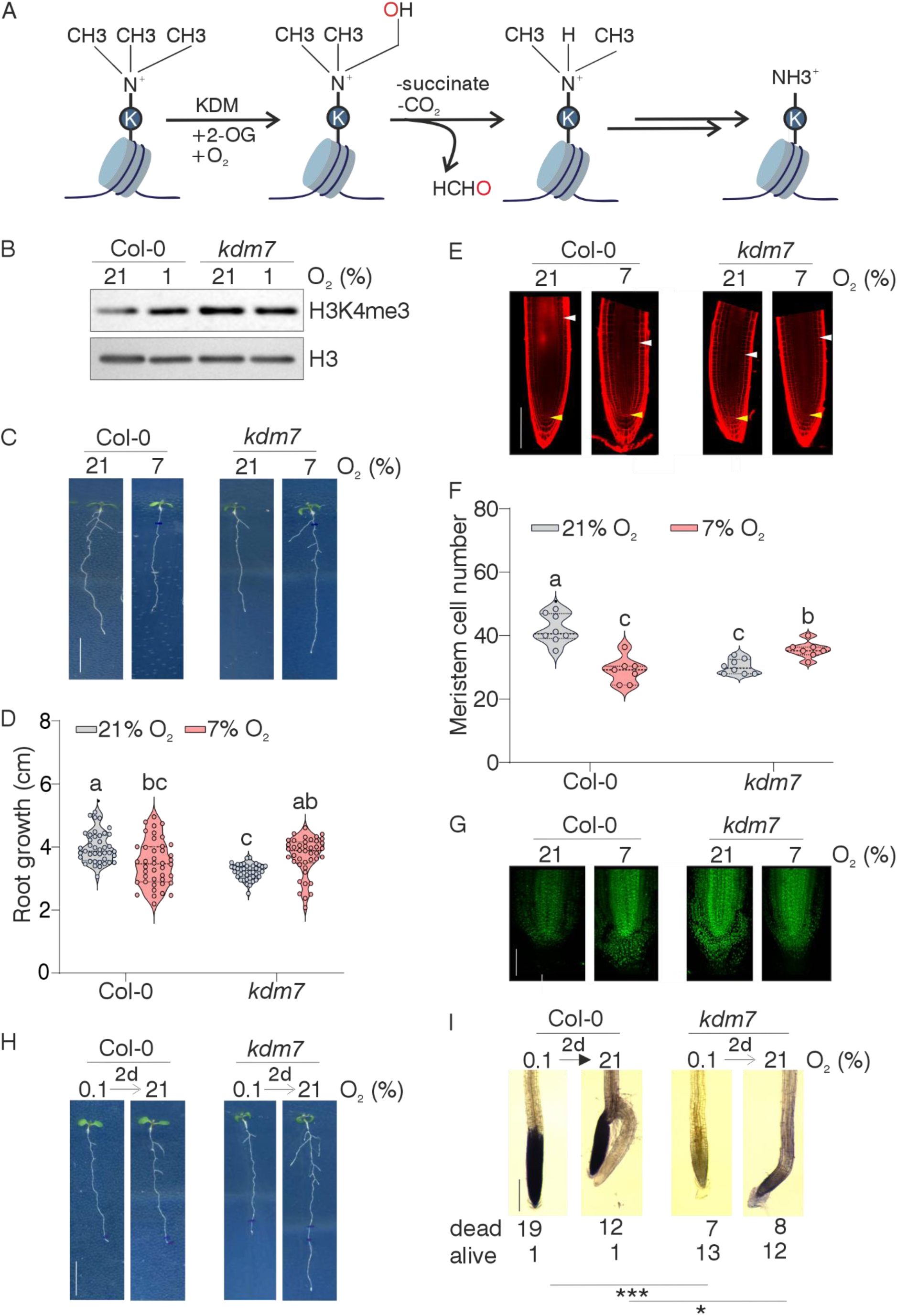
Aerobic H3K4me3 demethylation is mediated by KDM7s, which regulate root tip growth and hypoxia tolerance in *Arabidopsis*. (**A**) Schematic representation of lysine demethylation in histone tails by KDM enzymes. These enzymes use O₂ and 2-oxoglutarate (2-OG) as co-substrates to remove methyl groups, producing CO₂, succinate, and formaldehyde. (**B**) Detection of H3K4me3 after 12 h of normoxia (21% O₂) or hypoxia (1% O₂) in nuclear extracts from 20-day-old wild-type and *kdm7* plants. (**C–D**) Root length of wild-type and *kdm7* plants grown vertically for 7 days under normoxia (21% O₂) or mild hypoxia (7% O₂). (**E–F**) Root apical meristem length of wild-type and *kdm7* plants grown vertically for 7 days under normoxia (21% O₂) or mild hypoxia (7% O₂). (**G**) H3K4me3 immunostaining of wild-type and *kdm7* root tips exposed to mild hypoxia (7% O₂) or normoxia (21% O₂) for 12 h. (**H**) Post-hypoxia root growth phenotypes of wild-type and *kdm7* plants. (**I**) Root cell viability detected by Evans blue staining in wild-type and *kdm7* plants immediately and 2 days after exposure to 0.1% O₂ for 24 h. Frequencies of observed phenotypes are shown below the photographs (*p* ≤ 0.05; \*\**p* ≤ 0.001; Fisher’s Exact Test).

Suspecting substantial gene redundancy within this group, we set out to generate high-order mutants by crossing single T-DNA insertion lines (**Supplementary Fig. 4A-B**). A quadruple *kdm7acde* knock-out (KO) mutant did not show major differences in terms of plant growth and development, nor at the transcriptome level (**Supplementary Fig. 4C-F, Supplementary Table S1**). Inactivating the residual *KDM7B* in this background caused early arrest in 25% seed development and therefore a homozygous quintuple T-DNA mutant could not be obtained (**Supplementary Fig. 5A**). We resorted to expressing an artificial microRNA to silence *KDM7B* (**Supplementary Fig. 5B-C**).

Compared to the wild type, the resulting *kdm7* mutant exhibited increased H3K4me3 levels, which did not increase further under hypoxia (1% O_2_, 12 h) (**Fig. 2B**), suggesting that KDM7s play a major role in controlling overall H3K4 methylation levels under aerobic conditions. When grown aerobically on vertical plates for 7 days, *kdm7* mutants exhibited reduced root growth compared to the wild type, a condition that was fully reverted when plants where instead grown at 7% O_2_ (**Fig. 2C-D**). Microscopy of root apices revealed that hypoxia markedly reduced meristem size in wild-type plants but not in the *kdm7* mutant, where meristem cell number even increased compared with aerobic conditions (**Fig. 2E–F**). Immunostaining further confirmed elevated H3K4me3 levels in *kdm7* meristems, resembling those of hypoxic wild type (**Fig. 2G**). We next asked whether this increased H3 methylation state was sufficient to prime *kdm7* roots for survival under strict hypoxia, as mild hypoxia does in wild type (Fig. 1A–B). Indeed, *kdm7* root apices resumed elongation after 24 h at 0.1% O₂, whereas the same treatment killed the primary meristem in wild type (**Fig. 2H–I**). These findings suggest that mild hypoxia could prime roots to endure severe oxygen depletion by inhibiting KDM7 activity, thereby enhancing H3K4 trimethylation.

A 24 h mild hypoxia pretreatment prior to 24 h of strict hypoxia (0.1% O₂) promotes root growth resumption in wild-type *Arabidopsis thaliana* Col-0 and N-degron pathway mutants *ate1/2* and *prt6* (representative phenotype shown). (**C**) Quantification of post-hypoxia root growth. Roots exhibiting 0 ± 0.2 mm elongation were classified as “dead.” Mild hypoxia-pretreated (pink) and non-pretreated (grey) “alive” roots were compared using a t-test (*** p ≤ 0.001). (**D**) Detection of H3K4me3 through immunostaining after 3, 6, and 12 h of hypoxia (1% O₂) or normoxia (21% O₂) in total nuclear proteins extracted from 20-day-old *Arabidopsis* plants. (**E**) Effect of 1, 3, and 6 h of reoxygenation on H3K4me3 levels following 12 h hypoxia (1% O₂). (**F**) Detection of H3K4me3 after 0.5, 1, and 3 h of hypoxia (1% O₂) or normoxia (21% O₂) in nuclear extracts from 20-day-old *Arabidopsis* plants. (**G**) H3K4me3 detection after exposure of 20-day-old *Arabidopsis* plants to varying O₂ concentrations for 12 h. Total H3 is shown in D–G as a loading control. (**H**) H3K4me3 immunostaining of *Arabidopsis* root tips exposed to mild hypoxia (7% O₂) or normoxia (21% O₂) for 12 h.

### Hypoxic wild type and normoxic *kdm7* root tips show overlapping transcriptional profiles

To test our hypothesis, we compared the effects of genetic KDM7 inactivation with those of mild hypoxia (7% O₂, 7 days) in wild type using RNA-seq. We separately analysed root apices (first 5 mm from the tip), mature roots, and shoot apices from 7-day-old *kdm7* and Col-0 seedlings. KDM7 inactivation had a strong effect on root tip transcriptomes, but not on other tissues (**Fig. 3A, Supplementary Fig. 6**). Consistent with single-cell root atlas data (*22*), at least three KDM7s are expressed in the apical meristem: *KDM7A*, *B* and *D* (**Supplementary Fig. 7**). Interestingly, reduced KDM7 activity shifted the root tip transcriptome closer to that of mature roots (**Fig. 3A**). When examining the effect of mild hypoxia on the plant transcriptome, all three tissues showed a similar number of DEGs, although in root tips these were mainly upregulated rather than downregulated (**Fig. 3B, Supplementary Fig.6**), compatibly with a role of H3K4me3 in the activation of gene transcription (*23*). Notably, genes upregulated in *kdm7* root tips overlapped strongly with those induced by hypoxia in wild type apices (**Fig. 3C, Supplementary Table S2**), but only weakly with ERFVII-dependent hypoxia genes (**Supplementary Fig. 8**). The overlap between the transcriptome of hypoxic wild type root apices and aerobic *kdm7* ones was significantly enriched for genes involved in oxidative stress response, cell wall suberisation, glucosinolate biosynthesis and iron uptake regulation (**Fig. 3D** and **Supplementary Table S2**). Together with the results in **Fig. 2H–I**, these observations support a model where hypoxia-driven inhibition of KDM7 increases H3K4me3, activating genes required to sustain meristem activity under oxygen limitation.

**Figure 3.**
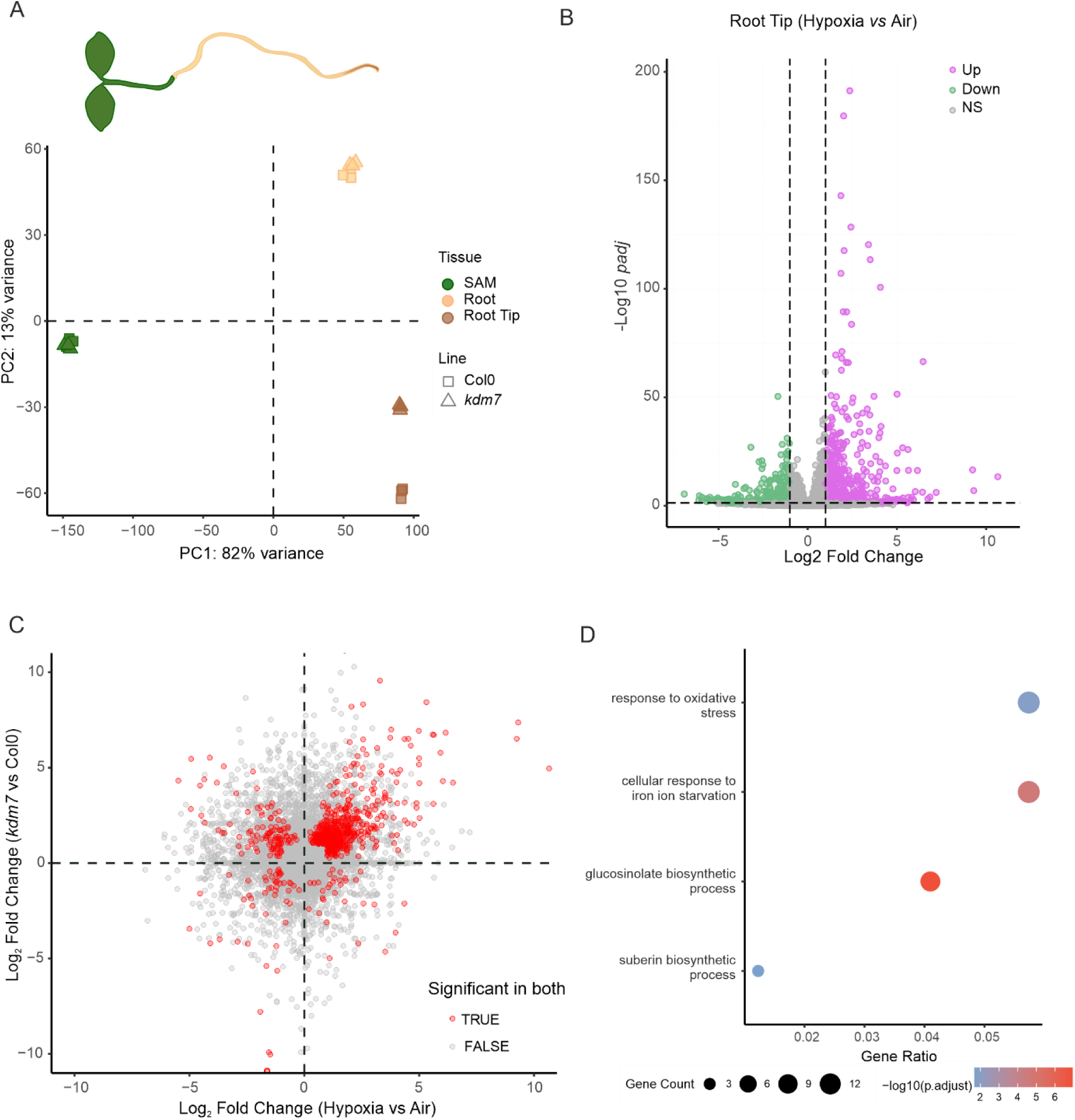
Genetic and hypoxic inactivation of KDM7 cause similar rearrangements of the root tip transcriptome. (**A**) Principal component analysis (PCA) of transcriptomes from shoots, root apices, and roots (excluding apices) of 1-week-old wild-type and *kdm7* seedlings. (**B**) Volcano plot showing the log₂ fold change (FC) in mRNA abundance between hypoxic (7% O₂) and normoxic (21% O₂) *Arabidopsis* root tips. (**C**) Scatter plot showing the relative changes in *Arabidopsis* gene expression under hypoxia (x-axis) and *kdm7* knockout (y-axis) conditions in root tips. Genes with significant log₂FC > |1| are shown in red. (**D**) Gene Ontology (GO) terms enriched among genes significantly induced in both hypoxic and *kdm7* root tips.

### H3K4 trimethylation in root tip cells activates iron deficiency signalling to increase iron uptake

Focusing our transcriptome analysis on iron signalling, we observed that group Ib basic helix–loop–helix (bHLH Ib) transcription factors and their FIT-dependent target genes involved in iron uptake were strongly upregulated in *kdm7* mutants (**Fig. 4A**). Upstream components of iron sensing and signaling, including positive regulators (group IVb/c bHLHs and the IRONMAN peptides IMA1 and IMA2) and negative regulators (BRUTUS (BTS) and BTS-like E3 ubiquitin ligases), were also induced, albeit more mildly (**Fig. 4A**). Consistent with these transcriptional changes, *kdm7* root tips accumulated significantly more iron than wild type (**Fig. 4B–C**), indicating impaired Fe sensing and enhanced uptake. We next asked whether elevated intracellular iron would support growth under conditions of extracellular iron sequestration. Indeed, *kdm7* mutants showed enhanced growth compared with wild type under iron-limiting conditions (**Fig. 4D–E** and **Supplementary Fig. 9C**).

**Figure 4.**
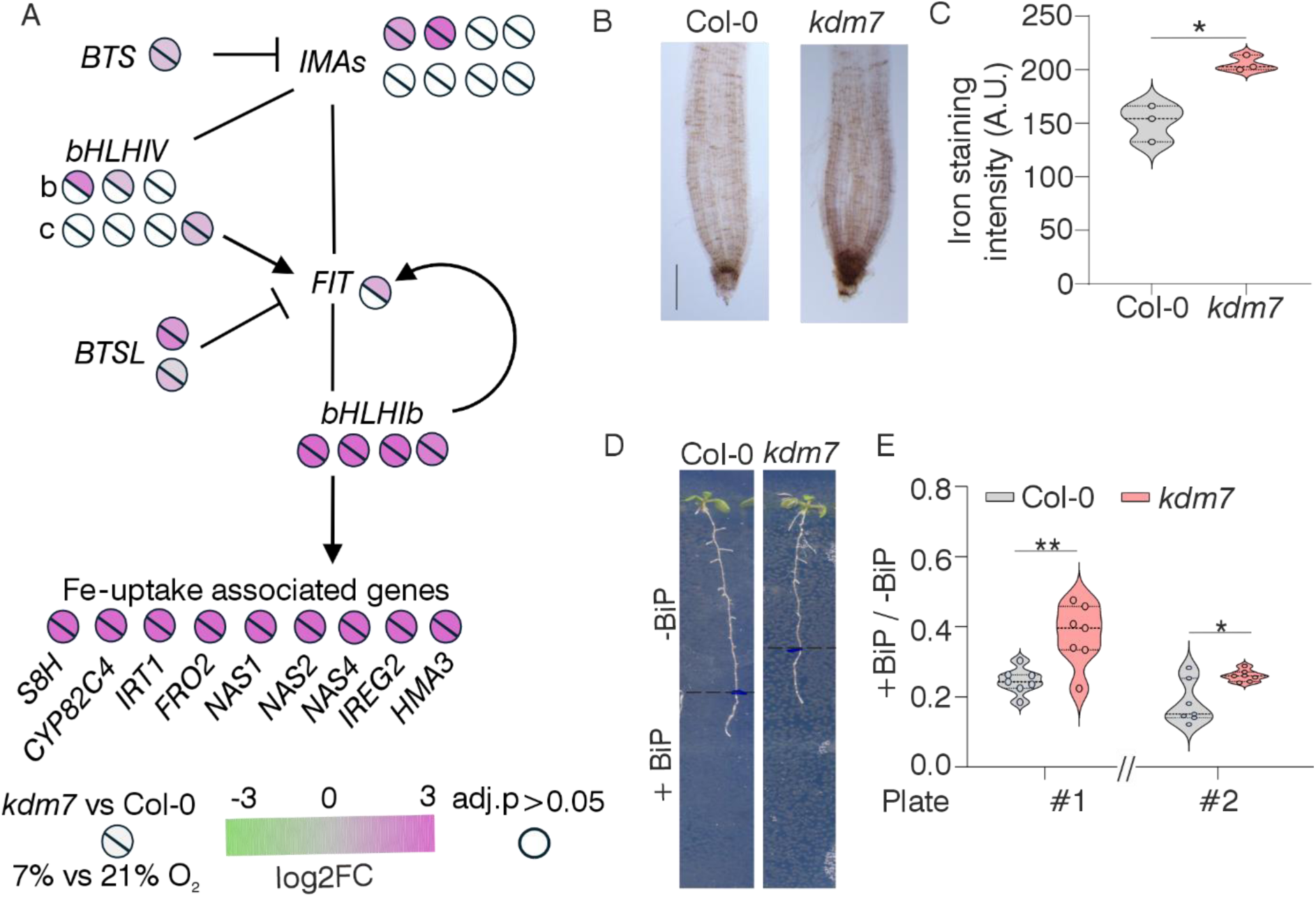
KDM7 inactivation induces the iron starvation response in root tips. (**A**) Heatmap showing the log2FC in expression of iron-signaling and iron-uptake genes in root tips following genetic (*kdm7* mutant) or hypoxic inactivation of *KDM7s*. (**B–C**) Iron accumulation in wild-type and *kdm7* root tips detected by Perls staining with DAB enhancement. (**D–E**) Root growth of wild-type and *kdm7* plants under bipyridyl (BiP)-induced iron starvation. The dashed lines in (**D**) indicate the root length at the time of transfer to BiP-containing medium. (**E**) Quantification of relative root length measured when plants were 5-days old and 2 days after transfer on 100 μM BiP medium. Statistical significance was assessed using a one-tailed *t*-test; *p* ≤ 0.05 (*), *p* ≤ 0.01 (**).

### KDM7s are directly inhibited by hypoxia

KDMs use oxygen and 2-OG as co-substrates to catalyse demethylation of lysine residues, with the release of CO_2_ and succinate as by-products (**Fig. 2A**) (*24*). Conversely, organic acids such as fumarate, succinate and 2-hydroxyglutarate (2-HG) have been shown to inhibit KDM activity **Supplementary Fig. 10A**) (*25*). Thus, hypoxia-induced inhibition of KDMs could result from either substrate scarcity or accumulation of inhibitors. Gas chromatography–mass spectrometry (GC-MS) analysis revealed no significant changes in 2-OG, 2-HG, fumarate, or succinate in seedlings exposed to hypoxia (1% O₂) for 3 h (**Supplementary Fig. 10B**). We therefore tested the alternative hypothesis that reduced oxygen levels in cells are sufficient to reduce KDM7 activity. To do this, we developed an in vivo H3K4 trimethylation reporter using NanoLuc Binary Technology (NanoBiT) based on split-nanoluciferase (nLuc) (*26*). We fused the N-terminal Large nLuc fragment (LgBiT) to the PHD domain of the TATA-box binding protein Associated Factor 3 (TAF3), a human histone reader specific for H3K4me3 (*27*), and separately created a chimeric version of Arabidopsis H3.1 whose N-terminal tail is extended with the C-terminal Small nLuc fragment (SmBiT). In cells expressing both constructs, a functional nLuc enzyme is expected to reconstitute and emit light in the presence of the necessary substrate when the N-terminal tail of the chimeric H3.1 protein is trimethylated at K4 (**Fig. 5A**). We confirmed that transiently transformed *Nicotiana benthamiana* leaf cells only emitted light when both components were expressed (**Supplementary Fig. 11A**). We also measured a significant increase in the signal when plants were exposed to hypoxia for 24 h (**Supplementary Fig. 11**).

**Figure 5.**
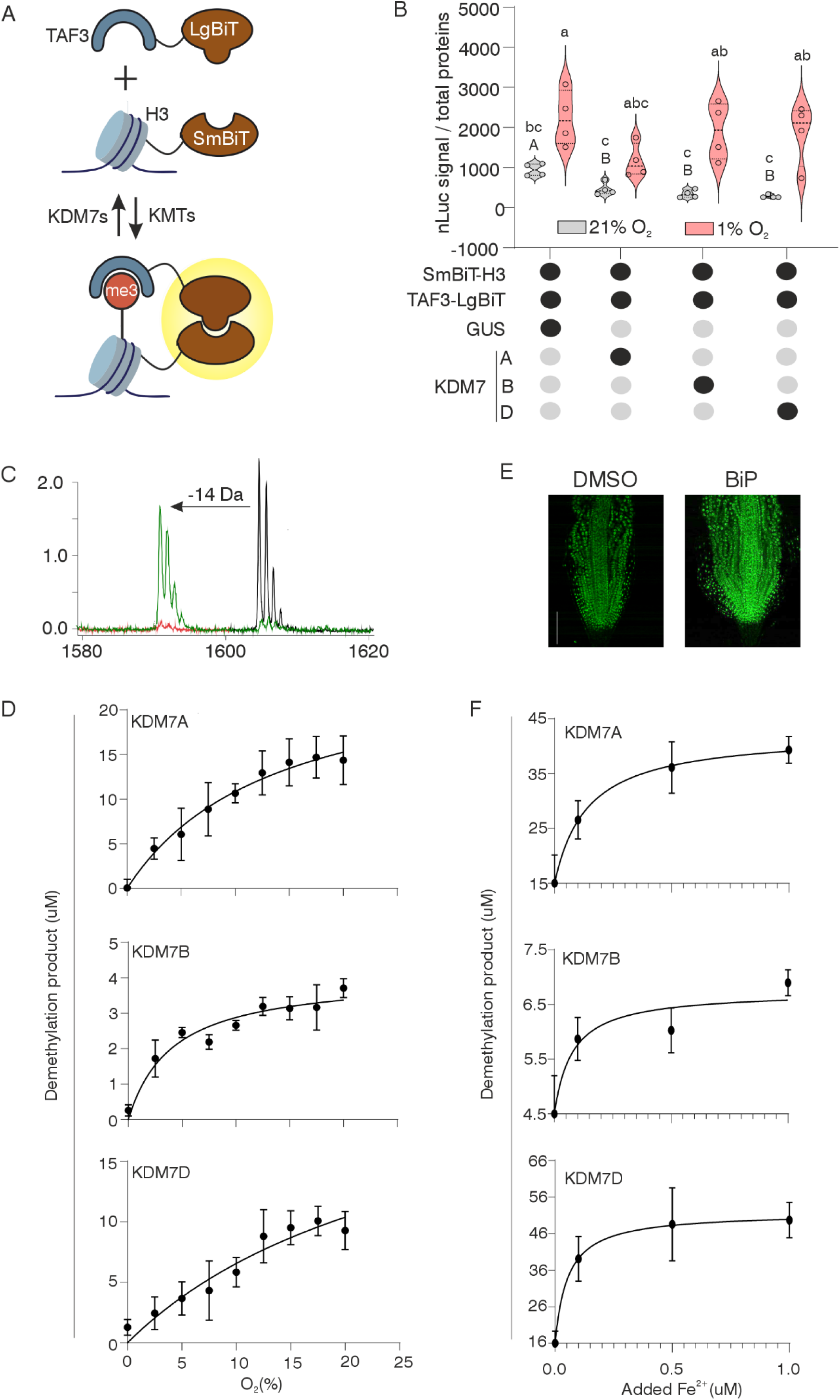
Low oxygen and iron-deprivation sensitivity of Arabidopsis KDM7s. (**A**) Schematic representation of the nanoBiT-based assay designed to test KDM activity in vivo: reconstitution of nLuc activity is mediated by binding of TAF3 to H3K4me3. (**B**) nLuc output of N. benthamiana leaf disks transiently transformed to express the reporters shown in A and KDM7A, B and D. β-glucuronidase (GUS) was used as a negative control. Different letters indicate statistically significant groups (p ≤ 0.05; two-way ANOVA for lowercase letters and one-way ANOVA for uppercase letters), followed by Tukey’s post-hoc test. (**C**) Example of mass spectra showing loss of 14 KDa in H3K4me3 peptides upon exposure to KDM7. (**D**) Demethylation activity of purified KDM7A, B and D at different oxygen concentrations. (**E**) Immonostaining of H3K3me3 in wild type root apices treated with 100 μM BiP. (**F**) Demethylation activity of purified KDM7A, B and D in reaction buffer with different concentration of Fe^2+^.

**Figure 6.**
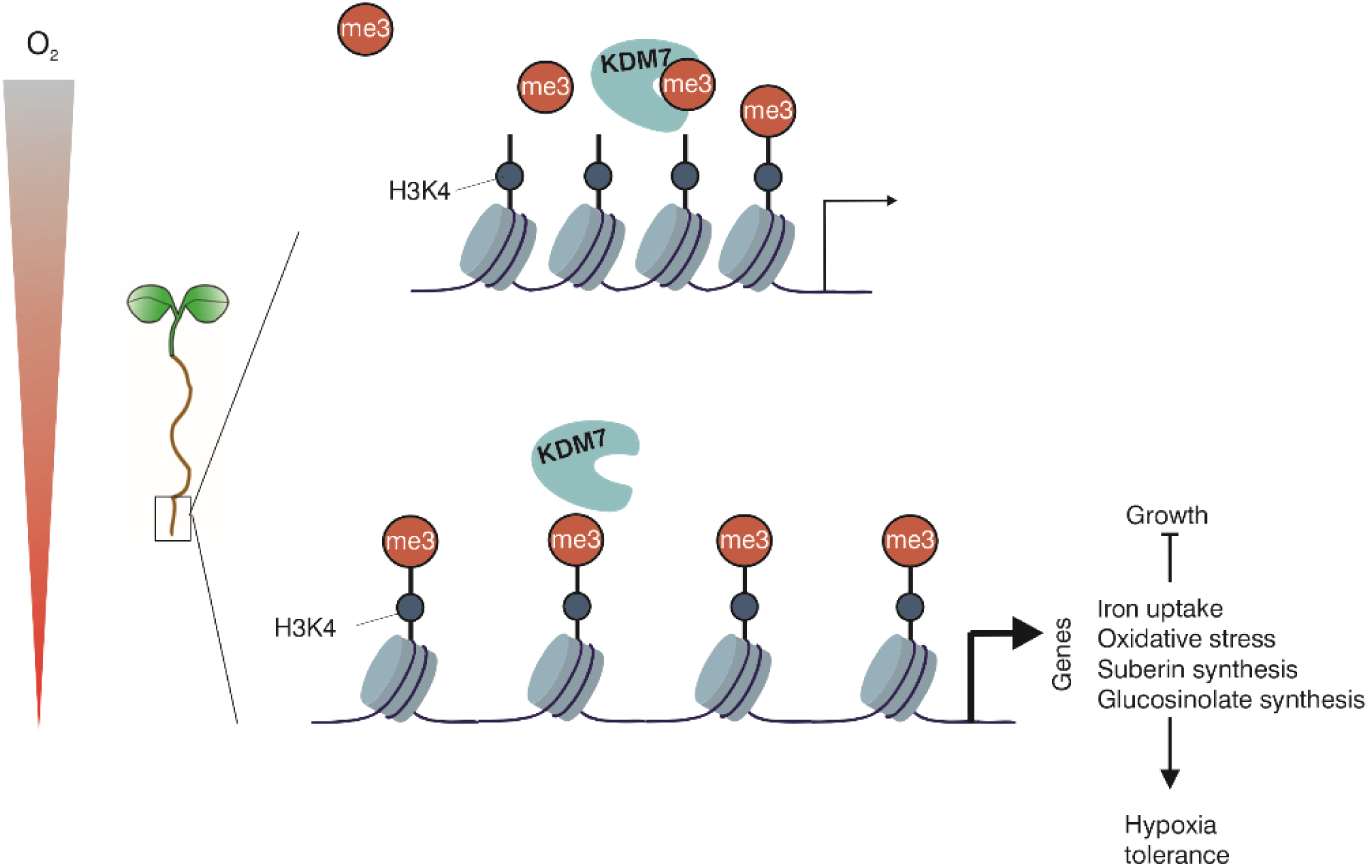
Proposed mechanism for hypoxia acclimation in roots. When root tips experience moderate hypoxia, KDM7 is inhibited, leading to increased H3K4me3. This leads to enhanced expression of iron starvation, oxidative stress and suberin synthesis-related genes, which prepare cells to endure severe hypoxia.

Having validated the effectiveness of our system to detect H3K4 trimethylation, we next tested the effect of overexpression of the three mostly expressed KDM7s in the Arabidopsis root meristem: KDM7A, B and D. Overexpression of all three KDMs reduced signal intensity compared with a beta-glucuronidase (GUS) control, compatibly with decreased trimethylation of H3K4 (**Fig. 5B**). Exposure to hypoxia (1% O2) for 24 h resulted in increased reporter signal, suggesting inhibition of the three KDMs (**Fig. 5B**). We next recombinantly expressed these three enzymes in insect cells, in order to test their sensitivity to hypoxia in vitro. Although suboptimal purification prevented us to determine precise enzyme kinetic parameters (**Supplementary Fig. 11B**), we were able to demonstrate that all three enzymes did indeed catalyse H3K4me3 demethylation at 21 % O_2_, while they displayed inhibition at reduced oxygen levels, notably in the range of physiological hypoxia (5-10%). This is consistent with the effects observed in vivo and indicates that the Arabidopsis KDM7s are sensitive to oxygen availability, potentially functioning as oxygen sensors, similar to the PCOs in the Cys-NDP.

Given the role of Fe²⁺ as a cofactor in KDM catalysis and the induction of iron-deficiency signaling and acquisition genes, we also assessed whether KDM7A, KDM7B and KDM7D could act as iron sensors too. The heterologously produced purified enzymes exhibited sustained activity at nanomolar Fe²⁺ ranges, suggesting that they are only affected by extreme iron scarcity in root cells (**Fig. 5E**). Moreover, immunodetection confirmed increased H3K4me3 under 2,2′-bipyridyl (BiP)-induced iron deficiency (**Fig. 5F**). However, these phenotypes are much milder than those observed for the Fe^2+^ sensors BRUTUS (BTS) and BTS-like (*28*), indicating that physiological iron limitation might not impact KDM activity and hypoxia is the main stimulus perceived by this epigenetic mechanism. The consequent increase in iron acquisition in root tips likely contributes to hypoxia priming by mitigating the oxidative stress elicited by hypoxia and subsequent reoxygenation (*29*).

## Discussion

In this study, we report the existence in *Arabidopsis* of an oxygen-sensitive system based on histone methylation that regulates adaptive responses at the epigenetic level in root apices. This system involves KDM7 enzymes, whose demethylase activity is impaired under mild hypoxia (7% O₂; **Fig. 2**), leading to increased H3K4me3 at chromatin regions encompassing genes involved in primary metabolism, oxidative stress tolerance, iron uptake, and glucosinolate biosynthesis (**Fig. 2–3**). The localized increase in H3K4 trimethylation promotes gene expression and primes root apical cells to withstand more severe hypoxic conditions (**Fig. 6**).

This resembles the epigenetic mechanism observed in human cells, where KDM4A, KDM5A and KDM6A inhibition by hypoxia causes increased H3K4, K9, K27 and K36 methylation that enables gene expression to control metabolic reprogramming and cell fates (*12*, *30*, *31*). This regulation was shown to establish the necessary chromatin landscape to facilitate the action of the master regulators Hypoxia Inducible Factors (HIFs) (*32*). In contrast, in the Arabidopsis root apex, we observed only modest overlap between ERFVII-regulated genes and those induced by KDM7 inactivation (**Supplementary Fig. 8**), suggesting that these systems operate in parallel rather than being strongly interdependent. Nonetheless, the observation that increased histone methylation via KDM inactivation contributes to hypoxia adaptation in both plants and animals underscores the evolutionary conservation of oxygen-sensing strategies in multicellular systems. Similar convergence has previously been noted for the role of the Cys-N-degron pathway (Cys-NDP) in controlling hypoxia responses (*33*).

While KDM7s’ role in enabling hypoxic priming is specific for root tips, our results do not exclude a similar role for other KDMs and increased methylation of other histone residues in the response to hypoxia in Arabidopsis. Our initial survey of histone methylation detected differences between aerobic and hypoxic seedlings, although these were not consistent over time, indicating an overlap with additional regulation, most likely linked to circadian rhythms (*34*). It will be interesting, in the future, to compare mutants in other classes of KDMs with the wild type for similar effects at the transcriptional level in other organs and/or tissues.

Our survey of histone methylation revealed substantial differences between hypoxic and normoxic whole seedlings. However, in fully differentiated root tissues and in the shoot, the overlap between genes differentially expressed under hypoxia in wild type and those altered in the *kdm7* mutant under normoxia was negligible (Fig. 3). This suggests that these tissues adjust their chromatin landscape to mount distinct responses, potentially through regulation of other histone-modifying enzymes. For example, increased H3K27 methylation by the PRC2 complex, stimulated by hypoxia-stabilized VRN2, was recently shown to regulate shoot growth (*17*, *35*).

The discovery of a novel oxygen-sensing mechanism in *Arabidopsis* raises the question of whether it is conserved in other angiosperms or more broadly across land plants. A recent survey of hypoxic responses revealed the absence of ERFVIIs in non-vascular species, while Cys2-ERFs, which are substrates of the Cys-NDP, are not stabilized under hypoxia (*2*). Given the similar hypoxia signalling pathway based on KDM inhibition observed in human and Arabidopsis cells it is plausible that epigenetic oxygen sensing predates ERF recruitment by the Cys-NDP. Future studies could address this hypothesis by examining the effects of hypoxia on histone methylation and comparing them to KDM inactivation in bryophytes.

In conclusion, our work identifies a tissue-specific oxygen-sensing mechanism based on KDM inactivation, which is essential in *Arabidopsis* for priming root acclimation to severe hypoxia (**Fig. 2**). This discovery opens the possibility of genetically or chemically manipulating KDM7s to enhance waterlogging tolerance in crops.

## Materials and Methods

### Plant material and growth conditions

*Arabidopsis thaliana* Columbia 0 (Col-0) was used as the wild ecotype throughout the whole study. The N-degron mutants *ate1/ate2* and *prt6-5* were the same as described in Licausi et al. (2011). The single T-DNA knock-out *kdm7* mutants were obtained from the Nottingham Arabidopsis Stock Centre (NASC): N655974 (Salk_135712) for KDM7B, N874918 for KDM7C, N874635 for KDM7D, N877462 for KDM7E and N688517 for *VRN2* respectively. Homozygous lines were identified via PCR screening of genomic DNA using gene-specific primers and T-DNA-specific primers listed in **Supplementary Table S3** as depicted in **Supplementary Figure 4**. High order mutants were obtained by crossing the single mutants and then screening the F_2_ generation as described above.

Soil grown plants were sowed on in moist peat soil containing pit and vermiculite in a 3:1 ratio, stratified at 4°C in the dark for 48 h and then germinated at 22°C day/18°C night with a photoperiod of 12 h of light and 12 h of darkness with 100±20 μmol photons m ^-2^ s^-1^ intensity. For plant growth under sterile conditions, seeds were surface sterilized with 70% ethanol (1 min), 0.85% sodium hypochlorite (10 min) and rinsed six times with sterile water. Sterile seeds were then sowed in half-strength Murashige–Skoog medium supplemented with 1% sucrose. Seeds were stratified for 72 h in the dark at 4°C and then transferred at 20°C with a 12-h light (100±20 μmol photons m ^-2^ s^-1^ intensity).

Hypoxia treatments were applied by mixing compressed air and pure N_2_ in a hypoxistation H35 (Don Whitley Scientific). All treatments were applied in the dark except for 7 day-long growth at 7% O_2_, which occurred under photoperiodic conditions (12 h light). Control plants were maintained at the same temperature and light conditions in an adjacent growth cabinet (Sanyo) located in the same room.

Growth in iron limiting conditions was performed as described by Grillet et al., 2018 (*36*). Briefly, Arabidopsis seeds were germinated in Estelle and Somerville (ES: 5 mM KNO₃, 2 mM MgSO₄, 2 mM Ca(NO₃)₂, 2.5 mM KH₂PO₄, 70 μM H₃BO₃, 14 μM MnCl₂, 1 μM ZnSO₄, 0.5 μM CuSO₄, 0.01 μM CoCl₂, and 0.2 μM Na₂MoO₄) media buffered at pH 5.5 with 1g l-1 2-(N-morpholino)ethanesulfonic acid (MES). The ES medium was either supplemented with 40 µM FeEDTA or without iron and 100 μM ferrozine to sequester traces of iron from media (0 µM Fe), or buffered at pH 7 with 1g l-1 3-(N-morpholino)propanesulfonic acid (MOPS) and supplemented with 10 µM FeCl_3_ (Non-available iron; NavFe). Seeds were stratified for two days and then grown under at 22°C with a 12-h light (100±20 μmol photons m ^-2^ s ^-1^ intensity). Root length was measured with FIJI (*37*) at 7 days post germination (dpg).

### CRISPR-mediated gene knock out

A single transgene bearing four guides directed against *KDM7A* (*AT2G38950/JMJ19*) (TGTGTTCAACCCAACCGAAG, CACATTCCTACACTTAGACG, ATGTGGGCGCTCCTAGAGTG and TGAACAGCAATATCCCCATG), spaced by a Gly-tRNA, was synthesised as flanked by the AtU3b promoter and a hepta-T PolIII terminator by GeneArt (Thermo Fisher Scientific). This DNA string was cloned into pDORN201 (Thermo-Fisher Scientific) and the resulting entry clone was recombined into pRU051 (*38*) using LR Clonase II (Thermo Fisher Scientific) following the manufacturer protocol. The resulting expression vector was transformed in *Agrobacterium tumefaciens* GV3101 by electroporation and the resulting transgenic bacteria used to transform *A. thaliana* Col-0 using the floral dip method (*39*). Transgenic seeds were selected based on green fluorescence and germinated in soil. The resulting plants were screened for the editing events that resulting in deletions of gene portions using primers KDM7B-g2F and KDM7B-g2R (**Supplementary Table S3**). T2 seeds were selected for homozygosity in KDM7 deletion and for the absence of the *Cas9* transgene.

### Artificial microRNA-mediated gene silencing

An artificial microRNA (amiRNA) against *KDM7B* (*AT4G20400/JMJ14*) was generated by overlapping PCR using the pRS300 vector as backbone (*40*) and subsequently cloned into pDORN201 (Invitrogen). The resulting entry vector was recombined into destination vector pO7WG2 using the LR reaction mix II (Invitrogen). The resulting expression vector was used for Agrobacterium-mediated transformation as described above for gene knock out. *KDM7B*-silenced plants were initially identified based on red seed fluorescence and subsequently by comparing *KDM7B* expression with wild type plants using primers sgKDM7Bfw and sgKDM7Brv (**Supplementary Table S3**).

### Baculoviral Protein Expression and Purification

Recombinant proteins KDM7A, KDM7B, and KDM7D were produced using the Bac-to-Bac™ Baculovirus Expression System (Thermo-Fisher Scientific) following the manufacturer’s protocol. Full-length coding sequences were cloned into a modified pACEBac1 vector (*41*) incorporating a Twin-Strep tag®, mScarlet fluorescent protein, and a 3C protease cleavage site **(Supplementary Table S4)**. Constructs were transformed into DH10Bac™ competent *Escherichia coli* (Invitrogen) for bacmid generation by site-specific transposition. Purified bacmid DNA was transfected into Sf9 insect cells to produce recombinant baculovirus stocks. Following infection, Sf9 cell pellets were washed with PBS, homogenized, and lysed in buffer containing 20 mM Tris pH 8.5, 200 mM NaCl, 4 mM DTT, 1 mM PMSF, 0.5% Tween, 0.1 mg/mL DNase I, 0.7 µg/mL Pepstatin, protease inhibitor cocktail, and 8.3 mM MgCl_2_. Cells were disrupted by sonication (Amplitude 40%, 15 s ON/15 s OFF cycles ×6) on ice. Lysates were clarified by centrifugation (4,000 rpm, 20 min, 4°C). Recombinant proteins were affinity-purified on Strep-Tactin resin (IBA LIfesciences), washed with 20 mM Tris pH 8.5, 200 mM NaCl, and 4 mM DTT, then eluted by 3C protease cleavage following overnight incubation at 4°C. Eluates were collected into 50 mL tubes with elution buffer (20 mM HEPES pH 7.5, 4 mM DTT). Purity was assessed by 10% SDS-PAGE under reducing conditions followed by Coomassie Blue staining.

### Histone lysine demethylation assay

Histone lysine demethylation assays were performed with modifications as described previously (*31*). Activity assays for KDM7 were conducted under the following conditions: 100 μM 2-oxoglutarate (2-OG), 100 μM L-ascorbate, 10 μM (NH_4_)_2_Fe(SO4)_2_·6H_2_O, and 10–350 μM H3K4me3 peptide (ARTK(me3)QTARKSTGGKA) in 50 mM HEPES buffer (pH 7.5). For oxygen-dependence assays, peptide substrate and buffer (87–92 μL) were equilibrated in septa-sealed glass vials at 25 °C for 5 min with controlled O_2_:N_2_ ratios using a mass flow controller. Subsequently, 1 μL each of 2-OG, L-ascorbate, and Fe^2+^ were added sequentially, followed by initiation of the reaction with 5–10 μL KDM7 enzyme via gastight Hamilton syringes. For the iron-dependence assay, (NH_4_)_2_Fe(SO4)_2_ was added at different concentrations as indicated in **Fig. 5F**. Reactions were incubated at 25 °C for various times and quenched with 900 μL 0.1% formic acid. Samples (1 μL) were analysed by MALDI-TOF mass spectrometry to monitor demethylation.

### Histone isolation

Histone isolated as described previously (*42*) with modifications. Two-week-old *Arabidopsis thaliana* seedlings (3 g) were ground in liquid nitrogen and resuspended in 10 ml of nuclear isolation buffer (15 mM PIPES pH 6.8, 5 mM MgCl₂, 60 mM KCl, 0.25 M sucrose, 15 mM NaCl, 1 mM CaCl₂, 0.8% Triton X-100, 1 mM PMSF, 0.7 µg/mL pepstain, and cOmplete™ EDTA-free protease inhibitor cocktail). The homogenate was filtered and centrifuged at 10,000 × g for 20min. Nuclei were extracted with 400 µL 0.4M sulfuric acid and histones precipitated with 132 µL acetone, then resuspended in 40 µL water.

### Western blotting

Histone extracts were dissolved in 2X Laemmli buffer and separated on a 10% polyacrylamide gel before transfer to a 0.2 µm PVDF membrane using Trans-Blot® Turbo™ Transfer System (Bio-Rad). Membranes were ‘blocked’ using PBS-T buffer (137 mM NaCl, 2.7 mM KCl, 10 mM Na_2_HPO_4_, 1.8 mM KH_2_PO_4_, 0.1% (w/v) Tween® 20) supplemented with 8g/L fat-free milk powder for 1 h at room temperature. Membranes were then incubated with appropriate dilutions of primary antibody in blocking buffer overnight at 4°C on an orbital shaker. The membrane was washed three times, 5 min each, with PBS-T before incubation with the recommended dilution of conjugated secondary antibody in blocking buffer at room temperature for 1 h. Following three washing steps (5 min each) with PBS-T, the membrane was incubated with luminescent substrate (West Dura, 34076, Thermo Fisher Scientific). Anti-H3K4me3 (Abcam, ab213224), anti-H3K9me3 (Insight Biotechnology, 230322/KG1/FD), anti-H3K27me3 (Insight Biotechnology, 230322/KG1/FD), anti-H3K36me3 (Insight Biotechnology, 230322/KG1/FD), anti-H3 (Abcam, ab209023), anti-H3K4me2 (Insight Biotechnology, 230322/KG1/FD), and anti-H3K4me (Insight Biotechnology, 230322/KG1/FD) were used at a 1:1,000 dilution as primary antibodies. An HRP-conjugated, goat anti-rabbit IgG H&L antibody from Abcam (ab205718) was used as the secondary antibody at a dilution 1:10,000. Signal was detected using the ChemiDOc XRS+ imaging system (BioRad). After immunoblot imaging, membranes were stained with Coomassie brilliant blue.

### Immunostaining and confocal microscopy

Immunostaining was performed following the established protocol described previously (*43*). Briefly, seven-day-old seedlings were fixed and treated with Driselase, followed by incubation with anti-H3K4me3 primary (Abcam, ab213224) and Alexa Fluor 488-conjugated (Invitrogen, A11008) secondary antibodies. Confocal images were acquired on a Zeiss LSM710 confocal microscope using 488 nm excitation and a 493–598 nm emission filter.

### Cell viability staining

Root tip cell viability was assessed using Evans blue staining following the protocol described previously (*44*). Briefly, 7-day old Arabidopsis seedlings were incubated in the staining solution (0.025% W/V Evans blue in double distilled water) for 15 min in the dark. Seedlings were washed three times with double distilled water and subsequently imaged using light microscopy (Zeiss LSM710, 10x objective).

### RNA isolation and qPCR

Fully developed leaves were harvested from soil-grown plants (4-week-old) and immediately frozen in liquid nitrogen. Total RNA, extracted as described by Kosmacz et al. (2015) (*45*), was subjected to DNase treatment using the RQ1-DNase kit (Promega). Five hundred ng total RNA were reverse transcribed into cDNA using the LunaScript® RT SuperMix Kit (NEB). Real-time PCR amplification was carried out with the QuantStudio™ 5 Real-Time PCR System (Thermo-Fisher Scientific), using qPCRBIO SyGreen® Mix (PCR Biosystems). *KDM7B* expression was measured using the *AT4G20400* specific primers and *Ubiquitin10* (*At4g05320*) was used as the housekeeping gene (**Supplementary Table S3**). Relative quantification of *KDM7B* expression of was performed using the 2^−ΔΔC(T)^ method (*46*).

### RNA isolation and RNA-sequencing

Total RNA was isolated from root tips (5 mm far from the root apex), developed roots (the remaining part of the root sample) and shoot apices (after removing developing leaves) of wild type Col-0 and *kdm7* (*abcde*) mutants. Whole 7-day old seedlings were used when comparing wild type and kdm7(*acde*). Total RNA was isolated from powdered frozen tissues using GeneJET Plant RNA Purification Kit (Thermo-Fisher Scientific). RNA sequencing was performed in paired-end mode (PE150) on a NovaSeq 6000 platform (Illumina) by Novogene. Raw sequence reads were quality-checked using FastQC v0.11.9 (*47*), and adapter sequences were trimmed with Trim Galore v0.6.5 (*48*). Clean reads were aligned to the *Arabidopsis thaliana* reference genome (TAIR10) using STAR v2.7.10 (*49*). Read counts per gene were obtained using featureCounts (Subread v2.0.4) (*50*). Differential expression analysis was performed in R v4.4.1 using the DESeq2 package (*51*) with the default Relative Log Expression (RLE) normalization method. Gene Ontology (GO) term enrichment analysis of differentially expressed genes (DEGs) was conducted using the clusterProfiler package v4.10.1.

### NanoBiT essay for histone lysine methylation

A sequence coding for SmBiT (VTGYRLFEEIL) fused to Arabidopsis H3.1 (At5g65360) was synthesized and cloned in an entry vector by GeneArt (Thermo-Fisher scientific) and recombined in pUBDEST (*52*) using LR clonase mix II (Thermo Fisher scientific). A second construct, coding for a fusion protein containing the SV40 nuclear localisation sequence, a codon-optimised 851-929 fragment of the human TAF3 and LgBiT was synthesized by GeneArt (Thermo-Fisher scientific) and cloned in pDONR201. The resulting entry vector was recombined in pF7WG2 - a customised plasmid generated by substituting the hygromycin resistance cassette with a green seed-fluorescent marker in pH7WG2 (*53*) - using LR clonase mix II (Thermo Fisher scientific). Coding sequences of *KDM7A* (*AT2G38950.1*), *KDM7B* (*AT4G20400.1*) and *KDM7D* (*AT1G08620.1*) were PCR amplified from cDNA and cloned in pDONR207. The resulting entry vector were recombined into pEARLYGATE202 (*KDM7A* and *KDM7B*) and pGWB514 (*KDM7D*) (*54*). All destination plasmids (**Supplementary Table S4**) were individually transformed into *A. tumefaciens* GV3101, which was then used to transiently transfect *Nicotiana benthamiana* leaves with single or multiple constructs, as described in **Fig. 5B** following the protocol described by Sparks et al. (2006) (*55*). Plants were incubated for 3 days under normoxia, then treated with normixa (21% O_2_) or hypoxia (1% O_2_) in the dark for 48 h. Leaf disks were excised from infiltrated regions and extracted in Passive Lysis Buffer (Promega). NanoLuc activity was measured with a Glomax® 20/20 luminometer (Promega) using the Nano-Glo® Luciferase Assay System (Promega), according to the manufacturer protocol.

### Cell iron staining

7-day-old seedlings were fixed and subjected to Perl staining, followed by signal intensification using 3,3’-diaminobenzidine (DAB), according to the protocol described previously (*56*).

### Metabolite Extraction and GC–MS Analysis

Two-week-old *Arabidopsis thaliana* plants (whole tissue; *n* = 8) were harvested following 3 h exposure to either 21% or 1% O₂ conditions. Immediately after collection, samples were snap-frozen in liquid nitrogen and stored at –80°C until extraction. Frozen tissues were homogenized using a Qiagen TissueLyser (Qiagen, Germany). Metabolites were extracted following a methanol–chloroform extraction protocol modified from Lisec *et al.* (2006) (*57*). Briefly, a known quantity of DL-norvaline (N7502-25G, Merck Life Science UK Limited) was added to each sample prior to extraction to serve as an internal standard for recovery monitoring and normalization. Homogenized tissue was mixed with extraction solvent and incubated in a Thermomixer (Eppendorf, Germany) at 70°C for 15 min with continuous shaking. The extract was clarified by centrifugation at 14,000 rpm for 10 min, and the resulting supernatant was transferred to a clean 2.0 mL microcentrifuge tube. Chloroform (250 µL) and water (500 µL) were added, and samples were centrifuged at 5,000 rpm for 15 min to achieve phase separation. From the upper polar phase, 100 µL was collected and freeze-dried under vacuum for GC–MS analysis. A parallel set of authentic metabolite standards, succinic acid (S3674-100G), fumaric acid (47910-5G), α-ketoglutaric acid disodium salt hydrate (K3752-5G), and L-α-hydroxyglutaric acid disodium salt (90790-10MG) (all from Merck Life Science UK Limited), was prepared and processed identically.

Dried extracts were derivatized prior to analysis. Each sample was resuspended in 25 µL of pyridine (EMSURE®, 1097280100, Supelco) and incubated in a Thermomixer for 30 min at 37°C and 900 rpm. Subsequently, 35 µL of *N*-tert-butyldimethylsilyl-*N*-methyltrifluoroacetamide (TBDMS, 375934-10X1ML, Merck Life Science UK Limited) was added, and samples were shaken for 30 min at 60°C and 900 rpm. GC–MS analysis was performed using an Agilent Intuvo 9000 GC system equipped with two 15 m DB-5ms Ultra Inert Intuvo GC capillary columns (0.25 mm ID, 0.25 µm film thickness; part number 122-5512UI-INT, Agilent Technologies LDA UK Limited) coupled to an Agilent 5977B mass spectrometer. A 1 µL injection was introduced in splitless mode. Mass spectra were acquired in selected ion monitoring (SIM) mode for the fragment ions *m/z* M–15, M–57, M–85, and M–159, corresponding to the derivatized forms of succinic acid-(TBDMS)₂, fumaric acid-(TBDMS)₂, α-ketoglutarate-(TBDMS)₃, 2-hydroxyglutaric acid-(TBDMS)₃, and the internal reference norvaline-(TBDMS)₂. Metabolite identification was based on comparison of retention indices and mass spectra with authentic standards and the NIST/EPA/NIH Mass Spectral Library (2011 version). Peak integration and deconvolution were performed using Agilent MassHunter Quantitative Analysis Software (revision B.08.00). Relative metabolite abundances were normalized to internal standard recovery and sample fresh (or dry) weight.

## Acknowledgments: Funding

This work was supported by the European Research Council (ERC) grant 101001320 (DaZ, VS and FL) and by the UKRI Biotechnology and Biological Science Research Council (BBSRC) grants BB/X001059/1 (FL and XC) and CBR01420 (FL and VS). SI was supported by a John Fell Fund (University of Oxford) grant.

## Author contributions

DaZ, IAS, EF and FL conceived the experiments, DaZ, SI, CE and YT provided the plant germplasm and insect cells, DaZ, ADC, PB and ADPB carried out the experiments, DaZ, FF, XC, AR, VS, FC and FMG analysed the data, BG, FMG, CE, DuZ, PRE, EF provided technical advice and analysis support. FL, DaZ and XC wrote the manuscript.

## Declaration of generative AI and AI-assisted technologies in the manuscript preparation process

During the preparation of this work, the authors used ChatGPT-5.1 to streamline the text and correct grammatical mistakes. After using this tool, the authors reviewed and edited the content as needed and take full responsibility for the final version of the published article.

## Data and materials availability

all data are available in the main manuscript and supplementary material. RNA-seq data are available at the NCBI SRA database (Bioproject PRJNA1346290).

## Supplementary Materials

### Supplementary Tables

**Supplementary Table S1.** Transcriptomic responses of Arabidopsis Col-0 and the kdm7acde quadruple mutant 7-days old seedlings to hypoxia.

**Supplementary Table S2.** Transcriptomic responses of Arabidopsis Col-0 roots, root tip, and shoot apical meristems (SAM) to mild hypoxia (7% O2) and to the kdm7 quintuple mutant in the same tissues.

**Supplementary Table S3.** Primer sequences used in this study.

**Supplementary Table S4.** List of plasmids generated in this study.

