## Supplementary Figures and Tables for "KDM7-mediated oxygen sensing reprograms chromatin to enhance hypoxia tolerance in the root"

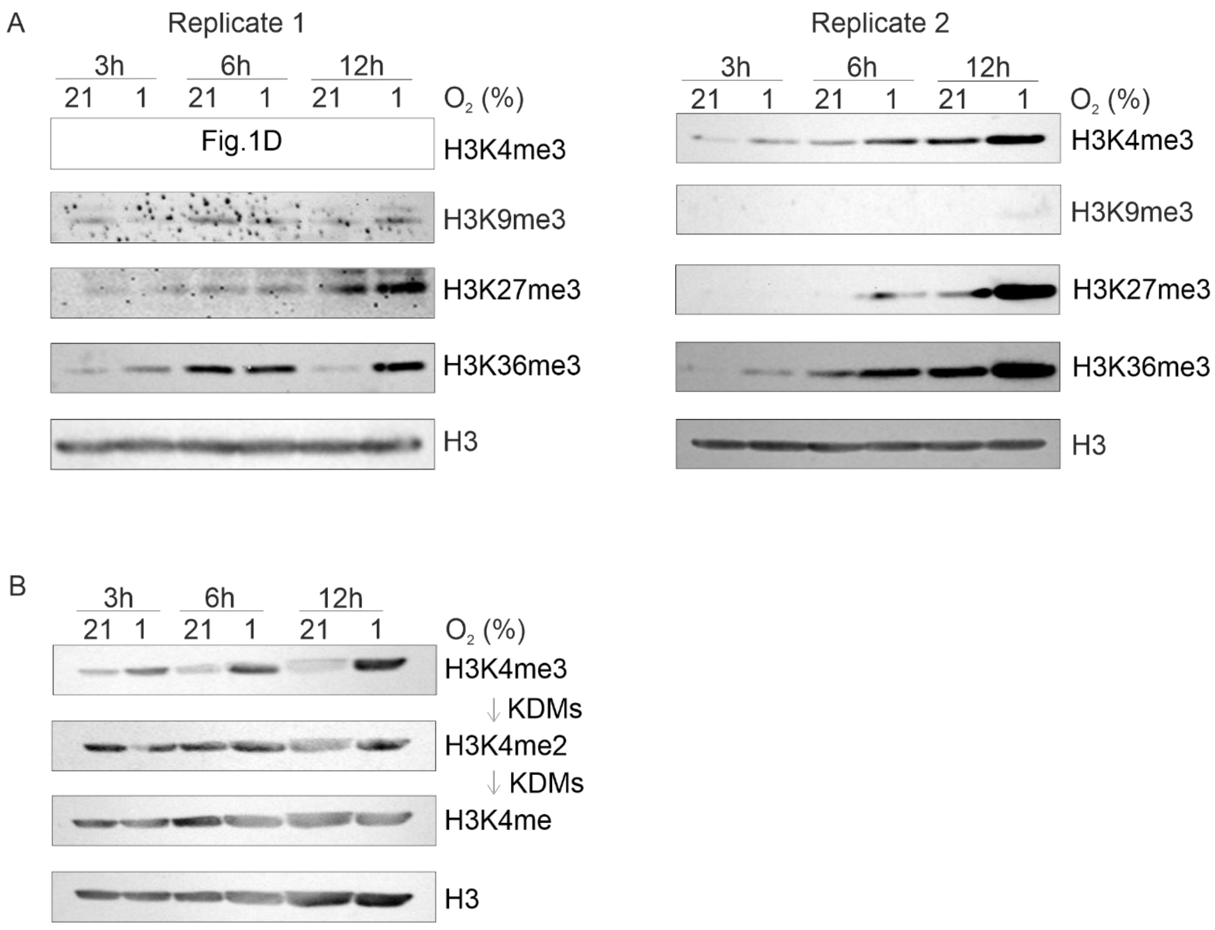


**Supplementary Figure 1. Effect of hypoxia on plant H3 methylation state.** (**A**) Quantification of lysine trimethylation in histone H3 (H3) from total protein extracts of 20-day-old *Arabidopsis* plants grown *in vitro* and exposed to hypoxia (1% $O_2$) for 3, 6, and 12 h. Methylation at K9, K27, and K36 was analyzed. H3K4me3 (replicate 1) is shown in Fig. 1D. Immunodetection of total H3 protein was used as a loading control. (**B**) Quantification of H3K4 trimethylation, dimethylation, and monomethylation in total protein extracts from 20-day-old *Arabidopsis* plants grown *in vitro* and exposed to hypoxia (1% $O_2$) for 3, 6, and 12 h. Immunodetection of total H3 protein was used as a loading control. KDMs are hypothesized to progressively remove methyl groups from H3K4.

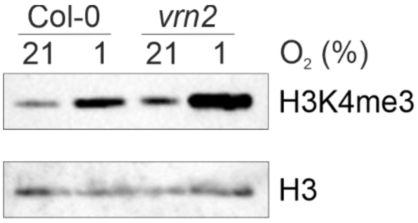


**Supplementary Figure 2. Effect of *vrn2* KO on Arabidopsis H3K4me3.** Quantification H3K4me3 from total protein extracts of 20-day-old *Arabidopsis* wild type and *vrn2* plants grown *in vitro* and exposed to hypoxia (1% $O_2$) for 12 h. Immunodetection of total H3 protein was used as a loading control.

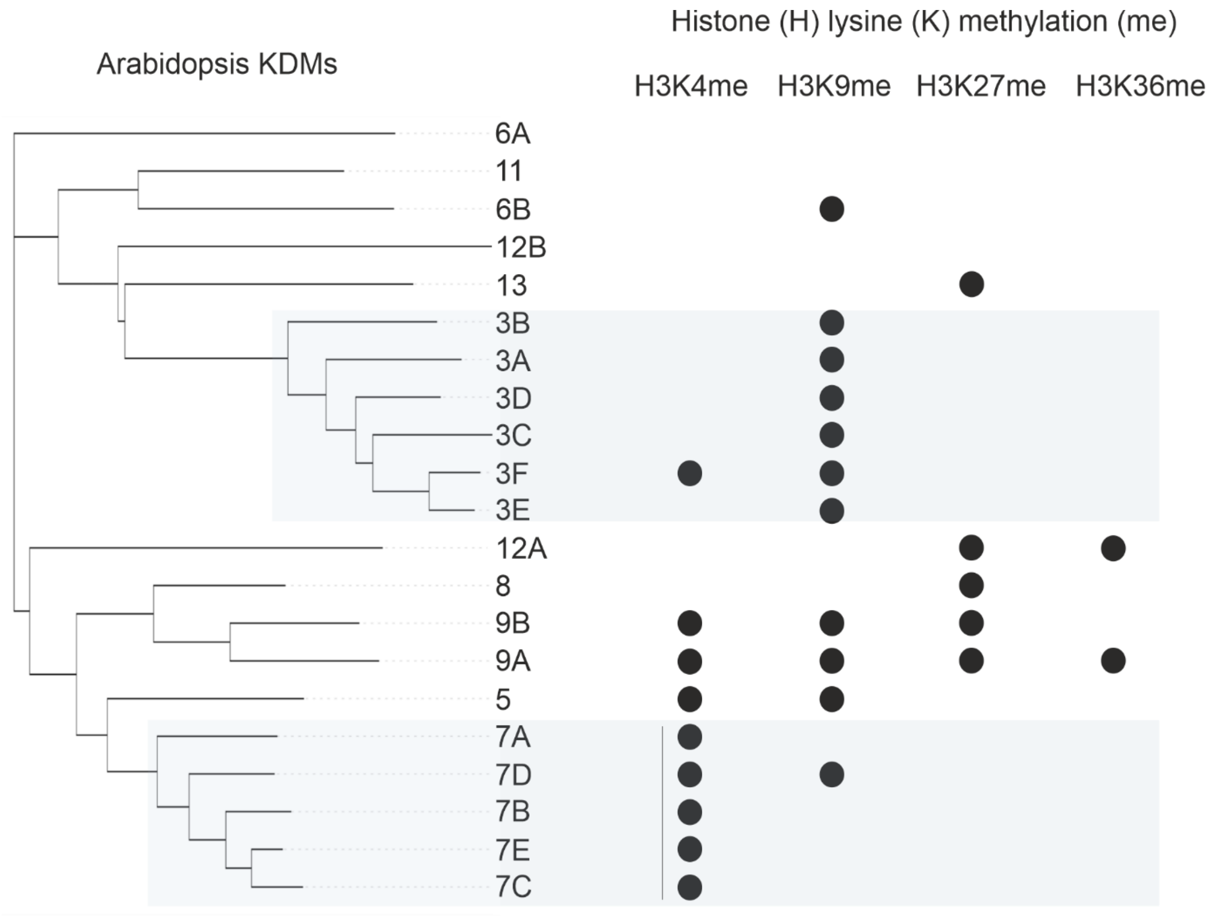


**Supplementary Figure 3. Unrooted phylogenetic tree showing the relatedness between KDM proteins in *A. thaliana*.** The activity on specific H3 lysine residues, as reported in the literature, is shown on the right. Relevant references for this are: https://theses.hal.science/tel-01124347v1/file/2014PA112093.pdf, https://doi.org/10.1104/pp.15.00520, 10.3390/genes12040529, 10.1111/tpj.13623, https://doi.org/10.1093/jxb/erz435 and 10.1111/j.1744-7909.2008.00692.x .

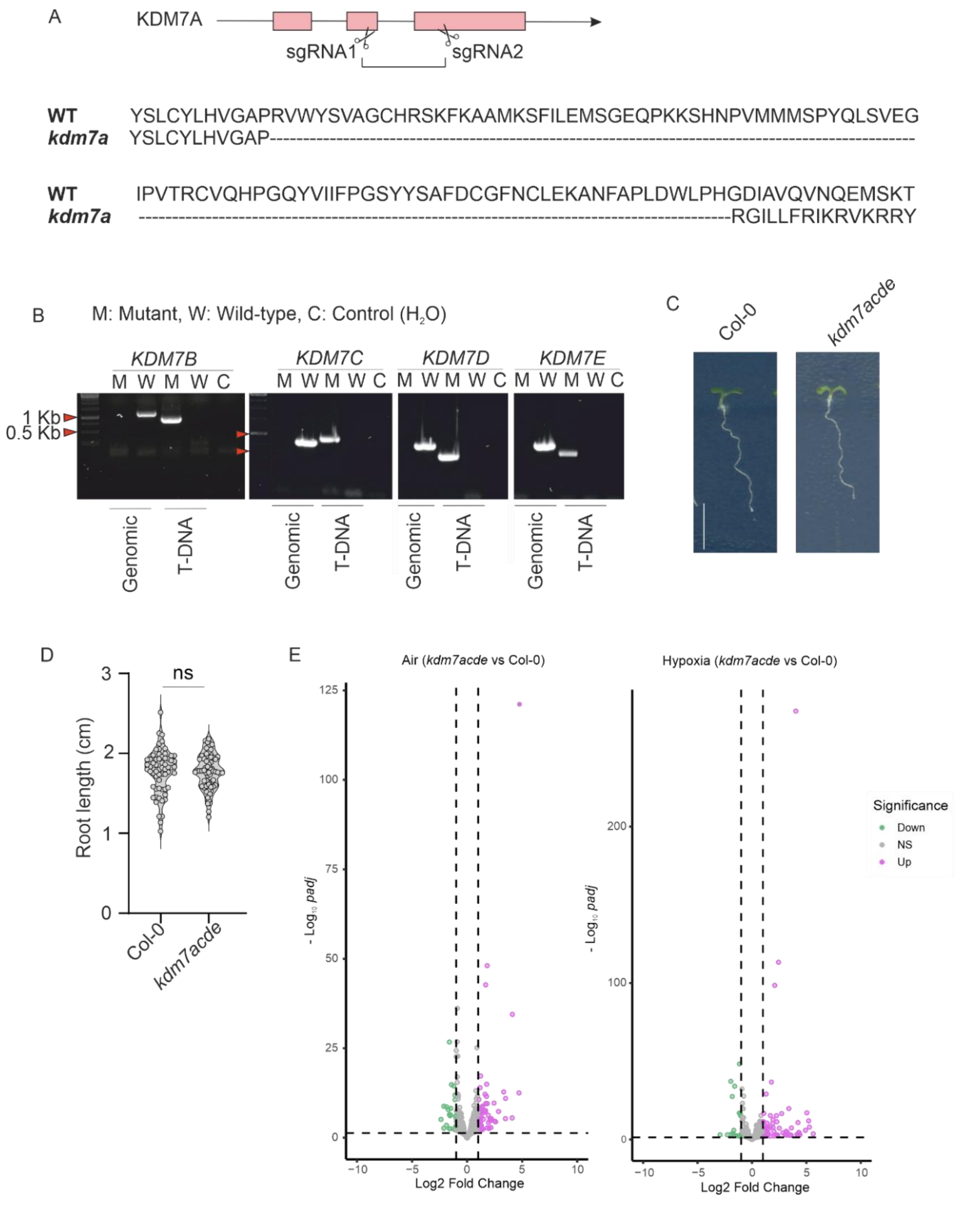


**Supplementary Figure 4. Generation of high-order Arabidopsis KDM7 mutants. (A)** CRISPR strategy adopted to knock out KDM7A. The portion of protein sequence deleted in this way, and the new sequence generated by frameshift, is also shown. **(B)** Screening of homozygous T-DNA insertion lines for *KDM7B* (N655974), *KDM7C* (N874918), *KDM7D* (N874635) and *KDMD7E* (N877462) using genomic and T-DNA primer combinations. M = mutant and W = wild type. (**C-D**) Comparison of wild type Col-0 and *kdm7acde* mutant phenotypes. D shows root length measurements. (**E**) Volcano plot showing the differentially expressed genes between the *kdm7acde* mutant and the wild type Col-0 under aerobic (21% $O_2$, left) and hypoxic (1% $O_2$) conditions.

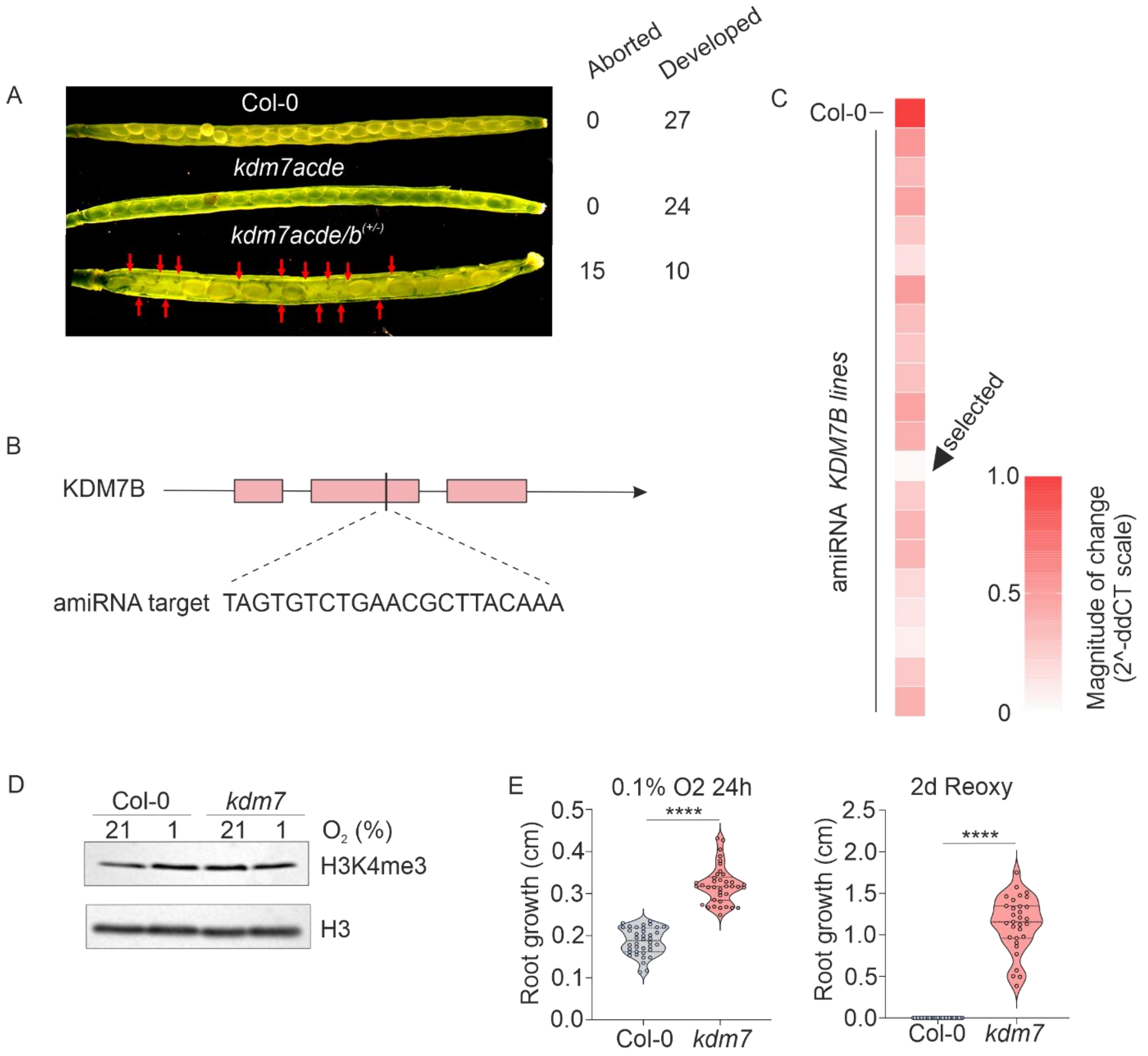


**Supplementary Figure 5. Generation and characterisation of a quintuple *kdm7* mutant.** (**A**) Seed abortion rates in Col-0, *kdm7acde* and *kdm7ab*$^{+/-}$*ced* mutants. (**B**) Artificial microRNA (amiRNA) targeting KDM7B. (**C**) Screening of *KDM7B*-silenced Arabidopsis plants. The selected line is shown with a black arrowhead. This line was named '*kdm7*' (**D**) H3K4me3 levels in wild type Col-0 and *kdm7* mutant. Total nuclear protein were extracted from 20-day-old *Arabidopsis* plants grown *in vitro* and exposed to hypoxia (1% $O_2$) for 12 h. Immunodetection of total H3 protein was used as a loading control. (**E** ) Primary root growth of wild type Col-0 and the *kdm7* mutant during 24h exposure to severe hypoxia (0.1% $O_2$) and 2 days after reoxygenation (21% $O_2$).

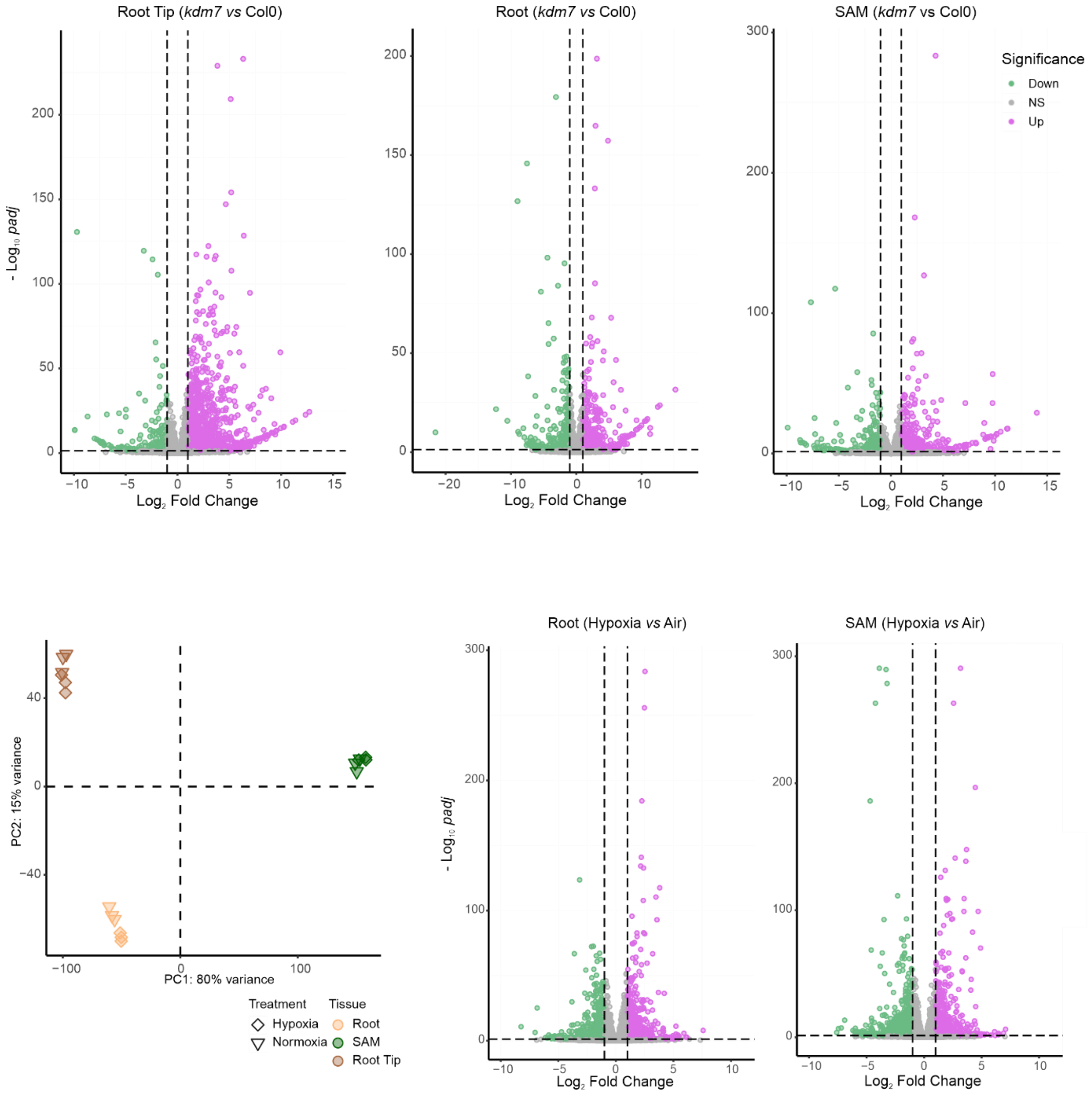


**Supplementary Figure 6. Transcriptome changes caused by KDM7 inhibition.** (A) Volcano plots showing $log_2$ fold change (FC) in mRNA abundance of *kdm7* mutants compared with wild type Col-0 in root apices, developed root tissues and shoots. (B) Principal component analysis (PCA) of transcriptomes from shoots, root apices, and roots (excluding apices) of 1-week-old wild-type seedlings. (C) Volcano plots showing $log_2$ fold change (FC) in mRNA abundance of developed roots and shoots from wild type plants grown in normoxia (21% $O_2$) and mild hypoxia (7% $O_2$) for 7 days.

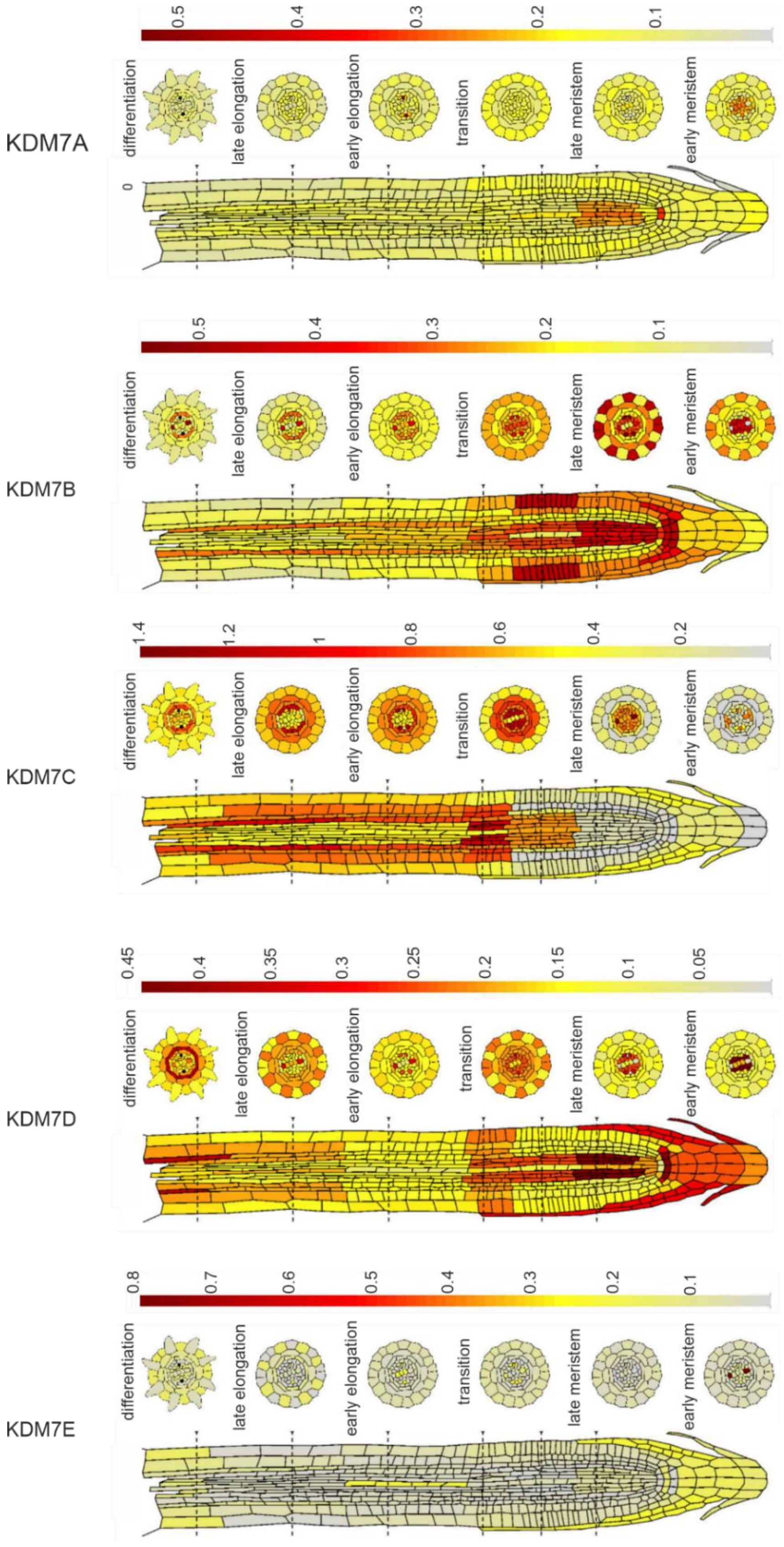


**Supplementary Figure 7. Expression atlas of KDM7 genes in the Arabidopsis roots.** Images were retrieved from the root cell atlas (https://rootcellatlas.org/).

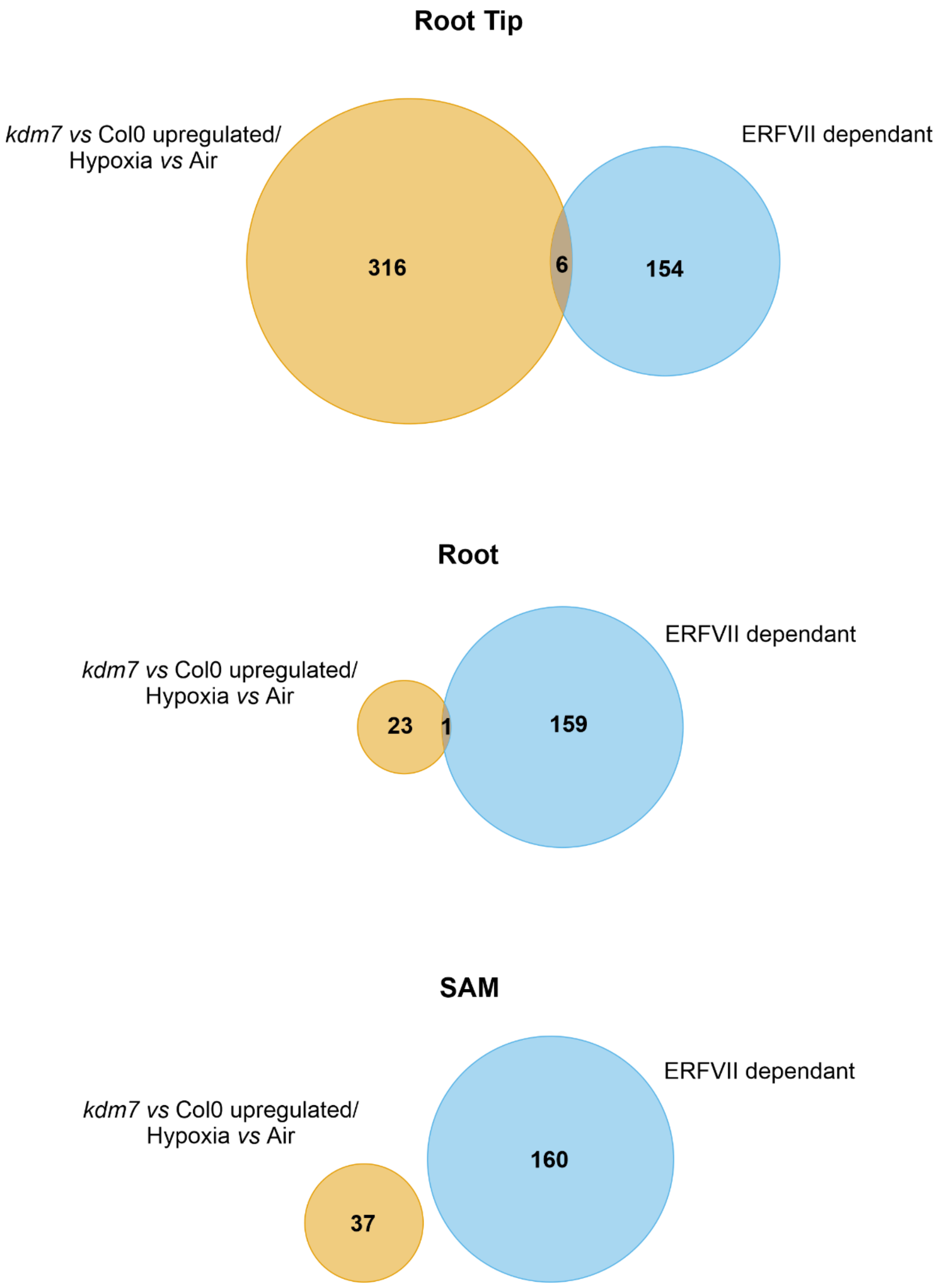


**Supplementary Fig. 8.** Venn-diagram showing the overlap between ERFVII-dependent hypoxia-induced genes and genes induced in *kdm7* mutants compared with the wild type Col-0.

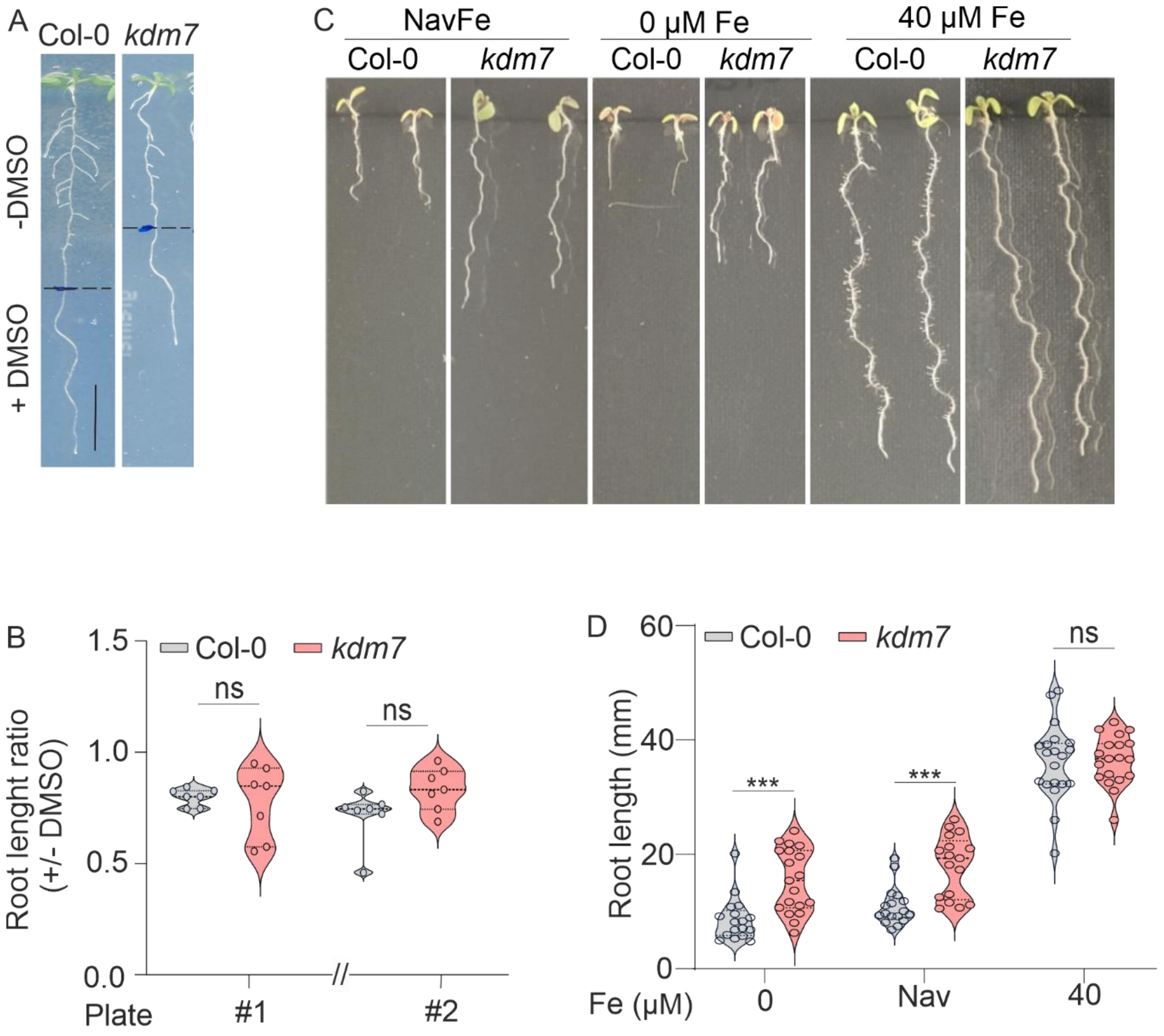


**Supplementary Figure 9. Effect of iron deprivation on root growth in Col-0 and the *kdm7* mutant. (A)** Root growth of wild-type and *kdm7* plants under bipyridyl (BiP)-induced iron starvation. The dashed line in indicates the root length at the time of transfer to BiP-containing medium. (**B**) Quantification of relative root length measured when plants were 5-days old and 2 days after transfer on BiP-free MS medium. (Absence of) statistical significance was assessed using a one-tailed t-test. (**C-D**) Primary root length of Col-0 and the *kdm7* mutant in media supplemented with an immobile iron source (non-available *iron*; *navFe*), no iron or 40 µM Fe2+.

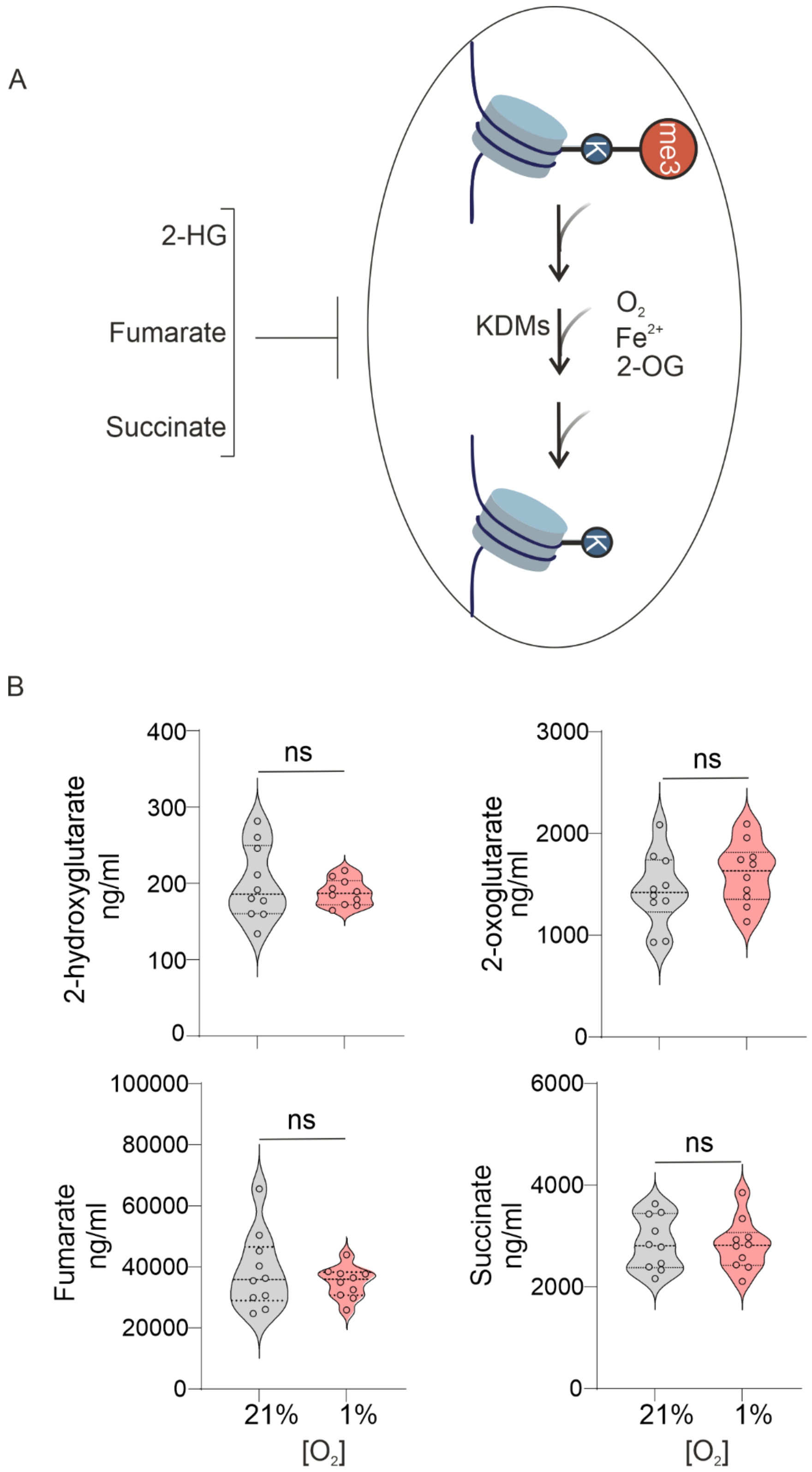


**Supplementary Fig. 10. Levels of organic acids whose abundance can potentially affect KDM activity in hypoxia.** (A) Schematic representation of the inhibition of histone demethylation by KDM caused by 2-HG, fumarate and succinate. (B) LC-MS quantification of 2-OG, 2-HG, fumarate and succinate in 20-day old seedlings treated for 3 h under normoxic (21% $O_2$) and hypoxic (1% $O_2$) conditions.

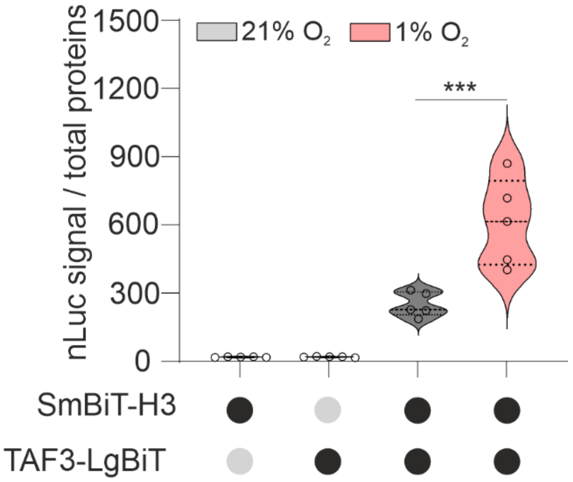


**Supplementary Fig. 11. Signal output for the NanoBiT-based reporter for H3K4me3.** Normalised nLuc activity in disks of *N. benthamiana* leaves infiltrated with the constructs LgBiT-TAF3 and SmBiT-H3 treated with hypoxia (1% $O_2$) and normoxia (21% $O_2$) for 24h.

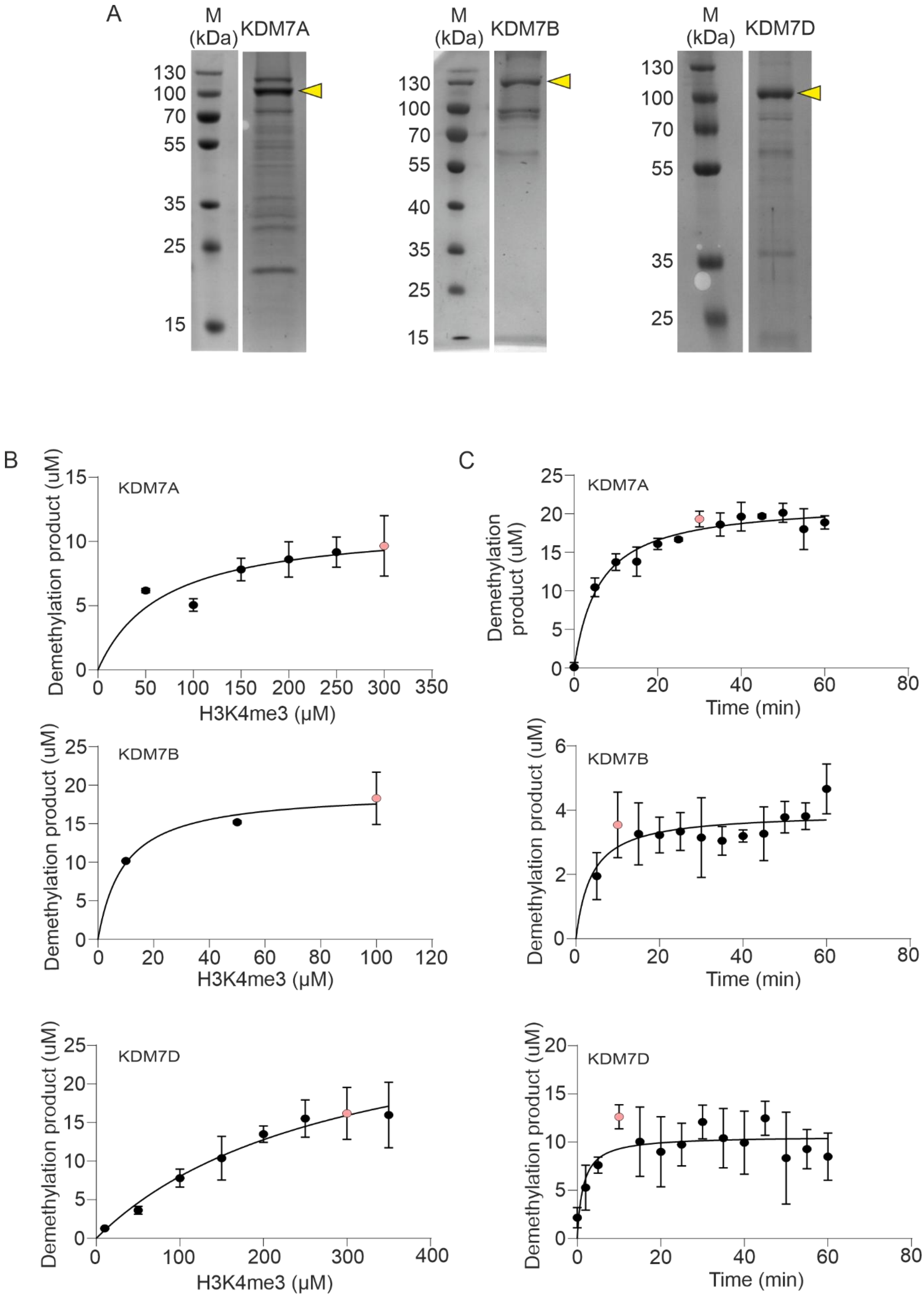


**Supplementary Figure 12. Purification of KDM7A, KDM7B, and KDM7D and their biochemical characterization.** (A) SDS–PAGE analysis of affinity-purified proteins following elution by 3C protease cleavage. The band corresponding to full-length KDM7 is indicated by a black arrowhead. (B) Titration of H3K4 peptide concentrations to determine saturating conditions for subsequent assays (highlighted by a pink dot). Reactions were incubated for 60 min. (C) Time-course analysis to determine the incubation period required for maximal accumulation of the demethylation product. The peptide substrate concentration used in this reaction corresponds to the one indicated by the pink dot in (B).
